# The actin-spectrin submembrane scaffold restricts endocytosis along proximal axons

**DOI:** 10.1101/2023.12.19.572337

**Authors:** Florian Wernert, Satish Moparthi, Jeanne Lainé, Gilles Moulay, Fanny Boroni-Rueda, Florence Pelletier, Nicolas Jullien, Sofia Benkhelifa-Ziyat, Marie-Jeanne Papandréou, Christophe Leterrier, Stéphane Vassilopoulos

## Abstract

Neuronal clathrin-mediated endocytosis has unique features in compartments such as dendrites and presynaptic boutons, but how membrane and extracellular components are internalized along the axon shaft remains poorly known. Here we focused on clathrin-coated structures and endocytosis along the axon initial segment (AIS), and their relationship to the periodic actin-spectrin scaffold that lines the axonal plasma membrane. Super-resolution optical microscopy, platinum replica electron microscopy, and their correlative combination on cultured hippocampal neurons reveal that in the AIS, clathrin-coated pits form on bare membrane patches, ∼300 nm circular areas devoid of spectrin mesh and lined by actin filaments we termed “clearings”. In fibroblasts and the proximal axon of neurons, spectrin depletion and drug-induced scaffold disorganization increase clathrin-coated pit formation. However, the presence of clathrin-coated pits at the AIS is not directly linked to actual endocytosis: using cargo uptake and live-cell imaging experiments, we find that most AIS clathrin-coated pits are long-lived and immobile within the spectrin mesh clearings. Direct perturbation of the spectrin scaffold as well as elevated neuronal activity could induce endocytosis downstream of clathrin pit formation, showing that spectrin clearings are structures responsible for regulated endocytosis at the AIS.

## Introduction

Endocytosis is a ubiquitous cellular process that allows cells to interact with their environment by internalizing surface receptors and extracellular substances (Mukherjee et al., 1997). Differentiated cells such as neurons extensively rely on endocytosis for retrieving and transporting membrane and extracellular components to organize their complex and compartmented morphology, and sustain the synaptic vesicle cycle at the core of neuronal communication. This role explains the historical focus on presynapses as the main site of endocytosis in neurons, with a wealth of studies dissecting its molecular mechanisms and coupling to the converse process of exocytosis (Azarnia Tehran and Maritzen, 2022; Rizzoli, 2014). Neuronal endocytosis has also been extensively studied in dendrites, where it drives receptor internalization near postsynapses (Rosendale et al., 2017; Catsburg et al., 2022) and is the source of a complex endocytic trafficking network (Yap and Winckler, 2022).

Outside of presynapses, endocytosis in neurons is mainly achieved through clathrin-mediated endocytosis (Camblor-Perujo and Kononenko, 2022). Clathrin forms triskelia made of trimerized heavy chains and light chains that assemble via adaptor proteins to form a cage-like coat around a membrane invagination called clathrin-coated pit (CCP), before scission of the CCP neck releases a clathrin-coated vesicle. In neurons, CCPs are found at the plasma membrane of the cell body and dendrites as well as in the vicinity of the active zone in presynapses, strengthening the prevalent view that endocytosis mainly occurs in these compartments. By contrast, little is known about the nanoscale organization or even the existence of clathrin-mediated endocytosis along the axon shaft and at the axon initial segment (AIS), the specialized compartment that constitutes the most proximal part of the axon (Rasband, 2010; Leterrier, 2018). It was commonly assumed that little to no endocytosis could take place along the axon shaft (Parton et al., 1992), a view reinforced by the presence of a dense submembrane assembly of spectrins, ankyrin G and anchored membrane proteins at the AIS, and the discovery of a dense, periodic scaffold of actin rings connected by spectrin tetramers lining the axonal plasma membrane (Xu et al., 2013; Lorenzo et al., 2023; Leterrier, 2021).

We previously used super-resolution microscopy combined with platinum-replica electron microscopy (PREM) on unroofed neurons to reveal the ultrastructure of the periodic actin-spectrin scaffold and AIS submembrane components (Vassilopoulos et al., 2019). Despite the presence of this dense undercoat, PREM views showed numerous CCPs present along the AIS of mature neurons, which was surprising because studies at the time only detected endocytosis in the nascent AIS in developing neurons (Torii et al., 2020; Fréal et al., 2019) or under acute excitotoxic stress (Benned-Jensen et al., 2016). More recently, robust endocytic activity at the AIS was proposed as a mechanism for membrane protein sorting, allowing the retrieval of somatodendritic proteins when they enter into the axon (Eichel et al., 2022). This prompted us to focus on the detailed architecture of CCPs at the AIS, their endocytic function and their relationships with the actin-spectrin submembrane scaffold.

We used a combination of super-resolution microscopy and PREM to visualize the structural components of the AIS and proximal axon at nanoscale resolution. We show that in addition to the periodic scaffold, AIS spectrins together with actin form well-defined circular exclusion zones we termed “clearings”, which allow CCPs to form on the bare plasma membrane area exposed at their center. Moreover, we demonstrate the role of the dense spectrin mesh in restricting the formation of these CCPs and their unusual stability, resulting in limited endocytosis of fluid phase markers at the AIS. Finally, we show that physiological stimulation of N-methyl-d-aspartate (NMDA) receptors releases this inhibition and triggers efficient endocytosis from AIS CCPs.

## Results

### Clathrin-coated pits are present in circular clearings of the actin-spectrin scaffold at the AIS

We first visualized the distribution of CCPs with regard to the spectrin mesh at the proximal axon of neurons after two weeks in culture labeled for α2-spectrin and for the AIS-specific adhesion molecule neurofascin (Figure 1, A). We used Structured Illumination Microscopy (SIM), which resolves the 190-nm periodicity of the actin-spectrin scaffold as well as individual CCPs, and found that CCPs were present in areas devoid of α2-spectrin, as confirmed by the intensity profiles of averaged images centered on CCPs (Figure 1, B). The presence of CCPs in areas devoid of spectrin is also observed further along the axon and similarly observed when labeling the spectrin mesh for ß2-spectrin (Supporting Figure 1, A-C). To isolate the axonal plasma membrane and associated cortical cytoskeleton in a close-to-native state, without additional signal from intracellular compartments, we mechanically unroofed cultured hippocampal neurons using ultrasound (Figure 1, C). The presence of CCPs inside circular areas devoid of α2-spectrin was made clearer by the unroofing, with a distinct circular hole of α2-spectrin around CCPs seen on the average image (Figure 1, D).

**Figure 1:**
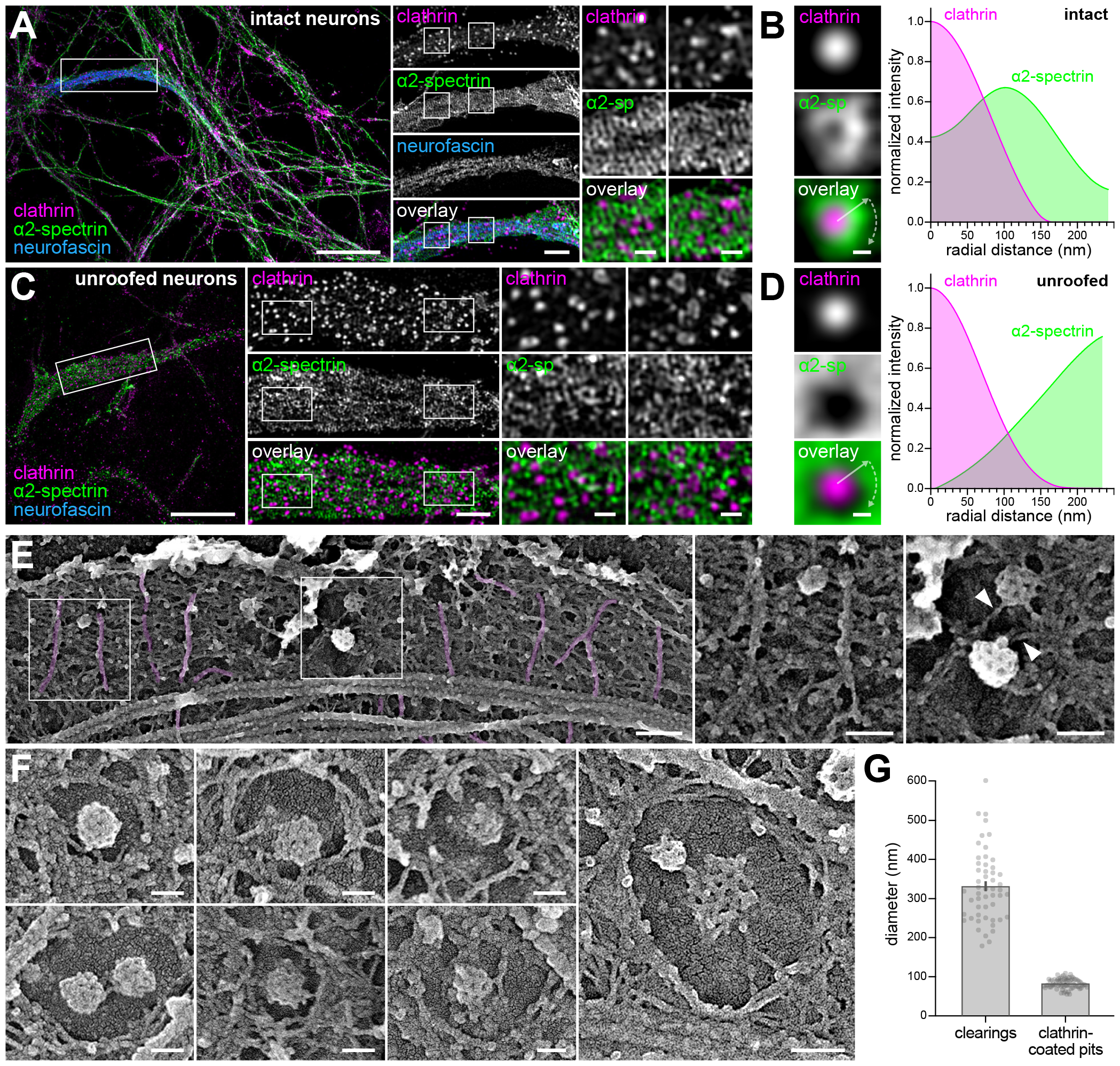
Clathrin-coated pits are present in clearings of the periodic spectrin scaffold at the AIS. A. SIM image of a cultured hippocampal neuron at 14 div fixed and stained for clathrin (magenta, CCPs appear as clusters), α2-spectrin (green, ∼190 nm-spaced bands are visible) and neurofascin (blue, labels the AIS). Scale bars, 10 µm (left image), 2 µm (center column), 0.5 µm (square zoomed images). B. Left, average images centered on CCPs for n=591 individual pits showing the distribution of clathrin (magenta on overlay) and α2-spectrin (green on overlay). Right, radial intensity profile corresponding to the average images on the left (gray arrow), showing how CCPs reside in an area devoid of α2-spectrin. Scale bar, 100 nm. C. SIM image of a neuron unroofed, fixed and stained for clathrin (magenta, CCPs appear as clusters), α2-spectrin (green, 190-nm bands are visible) and neurofascin (blue, labels the AIS). Central column shows zooms corresponding to the area highlighted in the left image, while square images on the right correspond to zooms on the areas highlighted in the central column. Scale bars, 10 µm (left image), 2 µm (center column), 0.5 µm (square zoomed images). D. Left, average images centered on CCPs for n=562 individual pits showing the distribution of clathrin (magenta on overlay) and α2-spectrin (green on overlay). Right, radial intensity profile corresponding to the average images on the left (gray arrow), showing how CCPs reside in an area devoid of α2-spectrin. Scale bar, 100 nm. E. PREM view of an unroofed proximal axon showing the presence of ∼190 nm-spaced actin rings (magenta), the dense mesh of spectrins between actin rings, and CCPs that reside in “clearings” of the spectrin mesh. Right images show zooms of square areas highlighted on the left image corresponding to an actin rings-spectrin mesh area, and an area with two CCPs (with an actin filament contacting the pit, arrowheads). Scale bars, 200 nm (left image), 100 nm (square zoomed images). F. Gallery of additional PREM views showing CCPs found in circular areas of bare plasma membrane devoid of spectrin mesh (“clearings”) along the proximal axon. Scale bars, 100 nm. G. Quantification of the diameter for spectrin mesh clearings (331 ± 13 nm, n=52) and for CCPs (83 ± 1.4 nm, n=75) from PREM views.

We then produced platinum replicas of unroofed hippocampal neurons to visualize the ultrastructure of CCPs at the AIS by PREM. To identify the axonal process stemming from unroofed cell bodies, we located the characteristic fascicles of AIS microtubules (Figure 1, E). High-magnification PREM views of the axonal membrane-associated cytoskeleton revealed the presence of characteristic actin rings embedded in a dense spectrin mesh as previously described (Vassilopoulos et al., 2019), but also provided unique views of CCPs at the AIS, with their characteristic honeycomb pattern (Figure 1, E). In agreement with super-resolution fluorescence microscopy, CCPs formed on a patch of bare plasma membrane, forming a circular “clearing” of the actin-spectrin scaffold. These clearings were devoid of proteinaceous material, save for single actin filaments which reached the central CCPs (Figure 1, F). Spectrin clearings measured 330 ± 12 nm in diameter, while the CCPs in their center measured 83 ± 1.5 nm (mean ± SEM; Figure 1, G). While single actin filaments were found to interact with CCPs in all neuronal compartments, we only observed the circular clearings of the actin-spectrin scaffold in proximal axons, while they were absent from cell body and dendrites that lack such a dense undercoat (Supporting Figure 1, C).

### Molecular organization of the CCPs and actin-spectrin clearings at the AIS

We next detailed the components and organization of the AIS CCPs and actin-spectrin clearings using Single Molecule Localization Microscopy (SMLM), alone and in correlation with PREM. 3D Stochastic Optical Reconstruction Microscopy (STORM) of clathrin in intact neurons revealed the presence of CCPs distributed circumferentially across the AIS (Figure 2, A-C). Point-Accumulation in Nanoscale Topography (PAINT) allowed us to simultaneously image CCPs and α2-spectrin in 3D at ∼20 nm resolution (Figure 2, D-E). This confirmed the presence of CCPs in areas devoid of the spectrin mesh, both on in-plane views and transverse sections of the AIS (Figure 2, F). After unroofing, CCPs were only seen along the remaining ventral plasma membrane by 3D-STORM (Supporting Figure 2, A-C). In addition, we used 3D-PAINT to show that the adaptor protein complex AP2 colocalizes to CCPs along the AIS (Supporting Figure 2, D-F), as classically described (McMahon and Boucrot, 2011; Pearse and Robinson, 1990).

**Figure 2:**
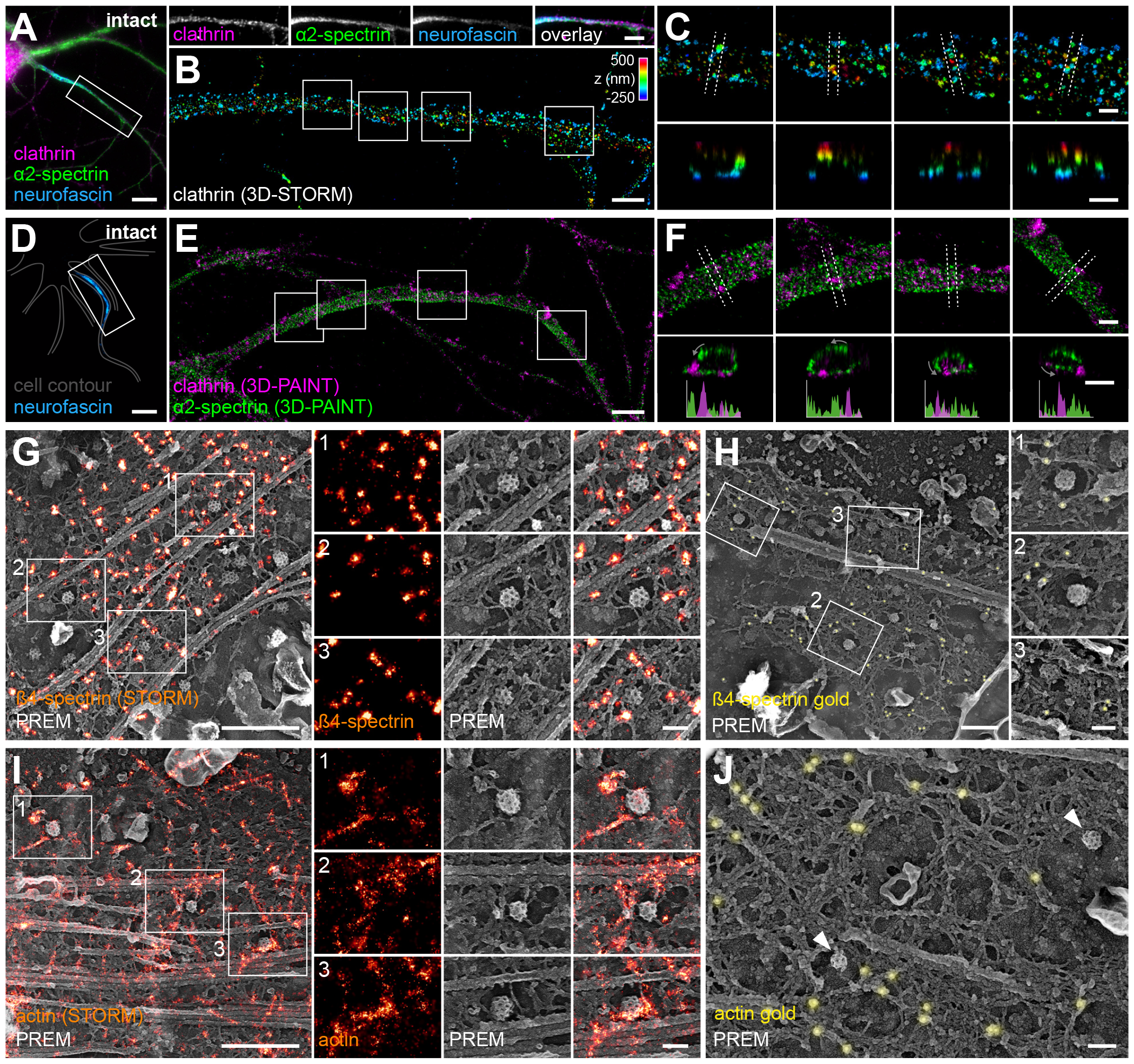
Molecular organization of the CCPs and actin-spectrin clearings at the AIS. A. Widefield image of a neuron fixed and stained for clathrin (magenta), α2-spectrin (green) and neurofascin (blue). Line of images on the right are individual channels and overlays corresponding to the AIS area highlighted on the left image. Scale bars, 10 µm (left image), 5 µm (AIS images). B. 3D-STORM image of clathrin corresponding to the AIS area shown in A, color coded for depth. Scale bar, 2 µm. C. Zooms showing individual CCPs on XY images (top row, corresponding to areas highlighted in B) and corresponding XZ transverse sections (bottom row, taken between the lines highlighted on the XY images). Pits are mostly found at the plasma membrane, delineating the axon contour on transverse sections. Scale bars, 0.5 µm. D. Widefield image of a neuron fixed and stained for neurofascin (blue). The contour of the whole neuron (gray) is delineated. Scale bar, 10 µm. E. 2-color 3D-PAINT images of the AIS corresponding to the image in A with staining for clathrin (magenta) and α2-spectrin (green). Scale bar, 2 µm. F. Zooms showing individual CCPs (magenta) in clearing of the periodic α2-spectrin mesh (green) on XY images (top row, corresponding to areas highlighted in E) and corresponding XZ transverse sections (bottom row, taken between the lines highlighted on the XY images). Pits are mostly found at interruptions of the spectrin mesh delineating the axonal plasma membrane on transverse sections, as shown by intensity profiles along the sections (bottom graphs). Scale bars, 0.5 µm. G. Correlative STORM-PREM image of an unroofed AIS labeled for ß4-spectrin (orange). Right, zoomed images showing CCPs in spectrin mesh clearings, with spectrin encircling the bare membrane area. Scale bars, 500 nm (left image), 100 nm (zoomed images). H. PREM view of an unroofed AIS immunogold-labeled for ß4-spectrin (yellow). Right, zoomed images showing CCPs in spectrin mesh clearings, with 15 nm gold beads bound to the spectrin mesh around the bare membrane area. Scale bars, 500 nm (left image), 100 nm (zoomed images). I. Correlative STORM-PREM image of an unroofed AIS labeled for actin (orange). Right, zoomed images showing CCPs in spectrin mesh clearings. Scale bars, 5 µm (left image), 100 nm (zoomed images). J. PREM view of an unroofed AIS immunogold-labeled for actin (yellow) showing CCPs in spectrin mesh clearings (arrowheads), with 15 nm gold beads bound to the spectrin mesh around the bare membrane area. Scale bar, 100 nm.

To further dissect the organization of the submembrane scaffold components around CCPs-containing clearings, we performed correlative STORM-PREM. After unroofing and fixation of neurons, we labeled a given component with Alexa Fluor 647 followed by STORM. We then produced platinum replicas of the same unroofed neurons and observed them by PREM (Vassilopoulos et al., 2019). We validated our correlative approach by using an anti-clathrin antibody to label CCPs at the AIS. The resulting overlays show how we can precisely correlate the 100-nm CCPs on both STORM and PREM images, although the anti-clathrin immunolabeling obscures the characteristic honeycomb pattern of the pits (Supporting Figure 2, G). We then stained the AIS-specific ß4-spectrin, localizing fluorescence clusters corresponding to the center of the spectrin tetramers within the dense mesh that surrounds CCP-containing clearings (Figure 2, G). We confirmed this localization using immunogold staining of ß4-spectrin, which similarly showed gold beads localized within the dense mesh (Figure 2, H). Turning to correlative STORM-PREM of actin, we observed fluorescent phalloidin delineating the 190 nm-spaced actin rings throughout the mesh, but also actin filaments along the circular border of the mesh clearings, as well as individual filaments contacting the CCPs (Fig.2, I). Indirect immunogold staining of actin filaments confirmed the presence of actin along the border of the clearings (Figure 2, J).

We also examined the nanoscale distribution of two more components of the actin-spectrin mesh in the AIS: the AIS master organizer ankyrin G (Rasband, 2010) and phospho-myosin light chain (pMLC) that decorates actin within the periodic scaffold (Berger et al., 2018; Costa et al., 2020). Correlative STORM-PREM and immunogold labeling showed that both proteins are present in the actin-spectrin mesh, including around the CCP-containing clearings (Supporting Figure 2, H-K). Altogether, our results indicate that in addition to forming a periodic actin-spectrin lattice, the AIS membrane-associated scaffold accommodates the presence of CCPs on the plasma membrane by forming stereotyped circular structures lined by actin filaments.

### The sparse submembrane actin-spectrin mesh restricts clathrin-coated structure formation in non-neuronal cells

Spectrin tetramers form hexagonal lattices in erythrocytes and organize periodically between actin rings in axons (Leterrier and Pullarkat, 2022). In non-neuronal cells such as fibroblasts, loose disordered spectrin networks are found co-existing with denser erythroid-like hexagonal and neuron-like periodic arrays (Ghisleni et al., 2020, 2023). We thus decided to first assess the role of actin and spectrins in regulating clathrin-mediated endocytosis in Rat2 fibroblasts, which express α2- and ß2-spectrin that can be targeted using the same interfering RNA sequences as in rat hippocampal neurons. PREM views of unroofed fibroblasts showed abundant clathrin-coated structures (CCSs) at their inner surface, with more diverse clathrin structures than in neurons (Figure 3, A), ranging from flat to fully curved endocytic pits (Heuser, 1980; Moulay et al., 2020). Using immunogold labeling of α2- and ß2-spectrin, we found both spectrins localized on the cortical mesh in the vicinity of CCSs (Figure 3, B). We also noticed that α2- and ß2-spectrin directly associated with actin filaments surrounding fully-formed CCPs (Figure 3, B). Correlative super-resolution spinning disk microscopy-PREM confirmed this observation and also showed that α2- and ß2-spectrin could form polygonal networks with actin filaments surrounding CCPs (Figure 3, C). We next used siRNAs to deplete spectrins, resulting in a ∼90% reduction of expression in fibroblasts (Supporting Figure 3, A-B) that did not affect the expression levels of clathrin heavy chain (Supporting Figure 3, C). PREM views of spectrin-depleted fibroblasts revealed a significant rise in the density of CCSs, which retained heterogeneous shapes (Figure 3, D): from 0.88 ± 0.09 CCS/µm^2^ in control cells to 1.8 ± 0.16, 1.6 ± 0.14, and 1.3 ± 0.14 CCS/µm^2^ for cells transfected with siRNA against α2-, ß2-, or α2- and ß2-spectrin, respectively (Figure 3, E). This suggests that the loose spectrin mesh found in non-neuronal cells can negatively regulate the formation of CCSs.

**Figure 3:**
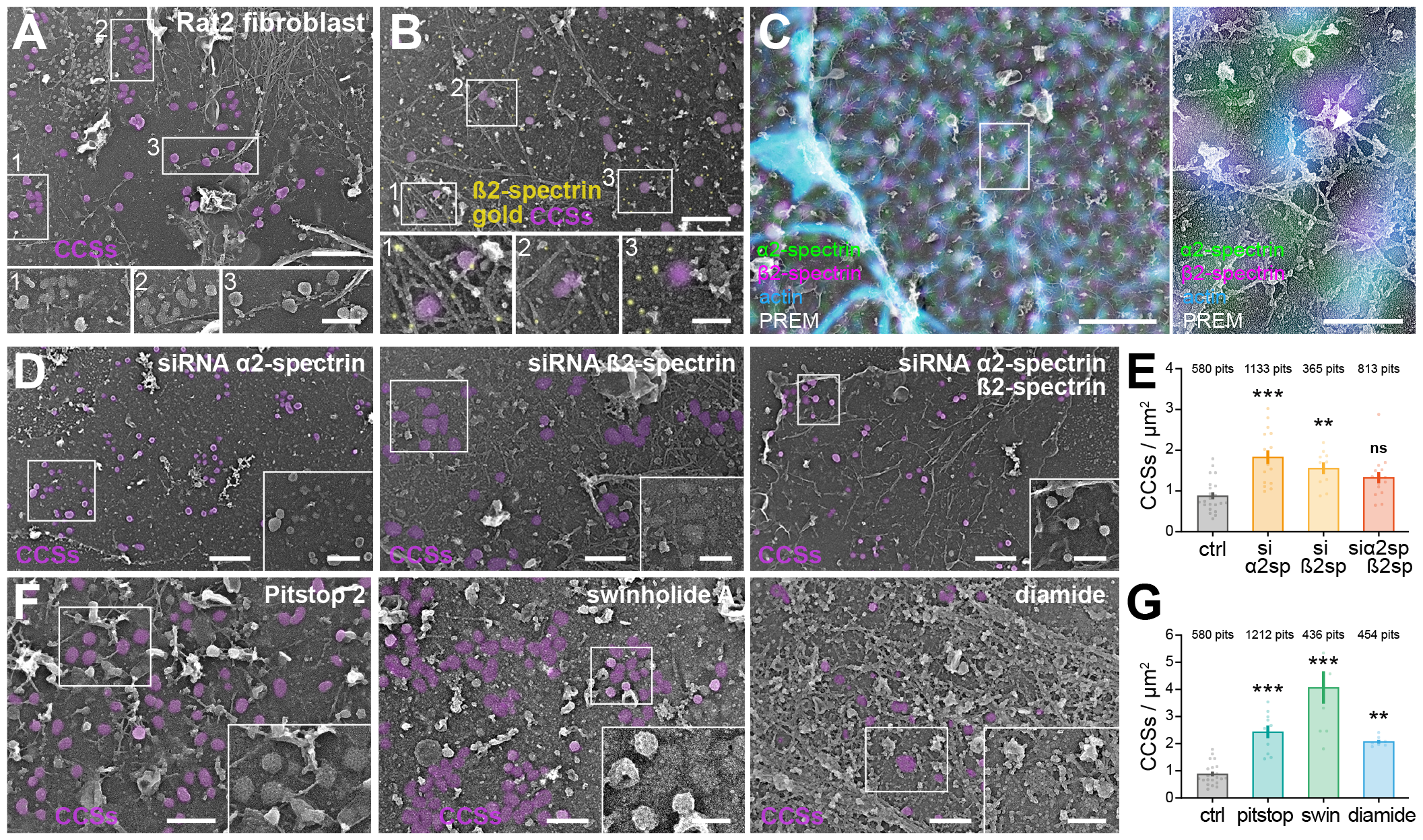
The sparse submembrane actin-spectrin mesh restricts clathrin-coated structure formation in non-neuronal cells. A. PREM view of an unroofed Rat2 fibroblast showing the presence of CCSs (magenta). Bottom images are zooms of the highlighted areas in the top image. Scale bars, 2 µm (top image), 200 nm (bottom images). B. PREM view of an unroofed Rat2 fibroblast immunogold-labeled for ß2-spectrin (yellow), with CCSs colored in magenta. Bottom images are zooms of the highlighted areas in the top image. Spectrins are present along the membrane, sometimes appearing as rods. Scale bars, 2 µm (top image), 200 nm (bottom images). C. Correlative super-resolution spinning disk microscopy-PREM of an unroofed fibroblast labeled α2-spectrin (green), ß2-spectrin (magenta), and actin (blue). Right images show a zoom of the area highlighted on the overlayed left image, with the spectrins being found along the areas rich in actin filaments, encasing a CCP (arrowhead). Scale bars, 1 µm (main image), 200 nm (right images). D. PREM views of unroofed Rat2 fibroblasts after knockdown of α2-spectrin (left), ß2-spectrin (center), or both (right), with CCSs colored in magenta on the main image. Zoomed inset corresponds to the area highlighted on the main image. Scale bars, 5 µm (main images for siRNA against α2-spectrin and α2/ß2-spectrin), 2 µm (main image for siRNA against ß2-spectrin), 200 nm (insets). E. Effect of spectrins knockdown on the density of CCSs as quantified from PREM views of unroofed Rat2 fibroblasts (n=25, 16, 10, and 15 images for Ctrl, siRNA against α2, ß2, and α2/ß2-spectrin conditions, respectively). F. PREM views of unroofed Rat2 fibroblasts after treatment with swinholide A (100 nM 3h, disassembles actin, left), pitstop2 (30 µM 15 min, inhibits endocytosis, center), or diamide (500 µM 15 min, perturbs spectrins, right), with CCSs colored in magenta on the main image. Zoomed inset corresponds to the area highlighted on the main image. Scale bars, 1 µm (main images), 200 nm (insets). G. Effect of drugs on the density of CCSs as quantified from PREM views of unroofed Rat2 fibroblasts (n=25, 9, 10, and 7 images for Ctrl, swinholide A, pitstop2, and diamide conditions, respectively).

We next used drugs to target CCSs, actin filaments, or the spectrin mesh and assessed their effect on CCS density in fibroblasts. We used Pitstop 2, a drug that selectively blocks endocytic ligand association with the clathrin terminal domain, to inhibit receptor-mediated endocytosis (von Kleist et al., 2011). Pitstop 2 treatment (30 µM, 15 min) led to a strong increase in CCS density to 2.5 ± 0.24 CCS/µm^2^ (Figure 3, F-G). To disassemble actin filaments including stable ones, we used swinholide A, which inhibits new filament assembly and disassembles existing ones (Vassilopoulos et al., 2019; Bubb et al., 1995). 100 nM swinholide A for 3h also caused a large increase in CCS density (to 4.1 ± 0.60 CCS/µm^2^) with clathrin lattices at all degrees of curvature occupying vast areas of the cell surface (Figure 3, F-G). We finally used diamide, a drug that causes oxidation of spectrins and disruption of their submembrane arrangement (Wu et al., 2017). PREM views on unroofed fibroblasts revealed that 500 µM diamide for 15 min causes a drastic alteration of the spectrin mesh, with aggregation of the submembrane scaffold into clumps (Figure 3, F). This disorganized spectrin mesh also caused the appearance of more CCSs, with a density rising to 2.1 ± 0.07 CCS/µm^2^ (Figure 3, G). Overall, perturbation of the actin-spectrin mesh increased the formation of CCSs at the membrane of non-neuronal cells.

### The periodic submembrane actin-spectrin mesh restricts CCP formation at the AIS

If the sparse actin-spectrin mesh can inhibit the formation of CCSs in fibroblasts, the denser periodic actin-spectrin mesh should be able to tightly control this process at the AIS of neurons. To test this hypothesis, we knocked down α2- and/ or ß4-spectrin in neuronal cultures using adeno-associated viruses (AAVs) expressing shRNA sequences. These AAVs could knock down spectrins efficiently, as shown by Western blots (Supporting Figure 3, E) and immunolabeling (Supporting Figure 4, A-D). In addition, AAV-expressed shRNAs against α2-spectrin, ß4-spectrin or a combination of both disorganized the periodic scaffold, as shown by the loss of periodic actin rings along the AIS and proximal axon (Supporting Figure 4, E-J). 3D-STORM showed that we could still unroof neurons depleted for both α2-spectrin and ß4-spectrin, revealing CCPs along the remaining ventral plasma membrane (Figure 4, A). PREM views of unroofed neurons depleted for α2- and/or ß4-spectrin showed a lack of submembrane spectrin mesh, with only isolated actin filament visible and the presence of numerous CCPs (Figure 4, B). Quantification confirmed that depletion of spectrins increased the density of CCPs in comparison with controls: from 1.9 ± 0.17 CCP/µm^2^ in control neurons to 6.4 ± 1.0, 5.7 ± 0.61, and 4.2 ± 0.87 CCP/ µm^2^ at the AIS of neurons depleted for α2-, ß4-, or α2- and ß4-spectrin, respectively; (Figure 4, C). This demonstrates that the dense spectrin mesh indeed restricts the formation of CCPs at the AIS plasma membrane.

**Figure 4:**
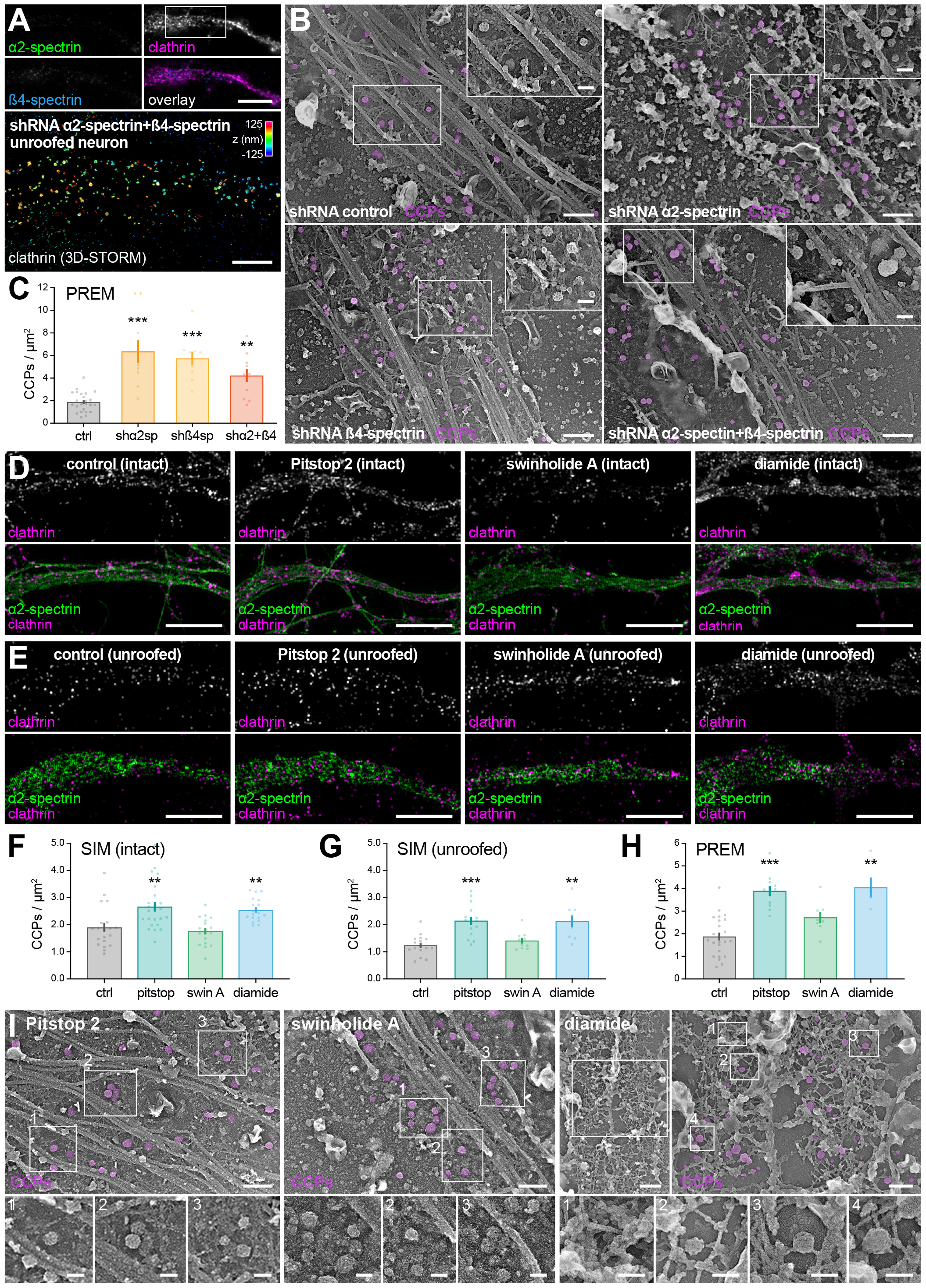
The periodic submembrane actin-spectrin mesh restricts CCP formation at the AIS. A. Top, widefield image of the AIS of a neuron infected with an AAV expressing shRNAs against α2-spectrin and ß4-spectrin, unroofed, fixed and stained for clathrin (magenta on overlay), α2-spectrin (green on overlay) and neurofascin (blue overlay). Bottom, 3D-STORM image of clathrin corresponding to the AIS area shown on the top image, color coded for depth. Scale bar, 10 µm (top images), 2 µm (bottom image). B. PREM views of the unroofed proximal axon of neurons infected with AAVs expressing control shRNAs (top left) or targeted against α2-spectrin (top right), ß4-spectrin (bottom left) or α2-spectrin and ß4-spectrin (bottom right), with CCPs colored in magenta on the main image. Insets are zooms of the area highlighted on the main images. Scale bars, 1 µm (main images), 200 nm (insets). C. Effect of spectrins knockdown on the CCP density as quantified from PREM views of unroofed proximal axons (n=25, 10, 10, and 11 images for Ctrl, shRNA against α2, ß4, and α2/ß4-spectrin conditions, respectively). D. SIM images of the proximal axon of neurons treated with vehicle (0.1% DMSO 3h), swinholide A (100 nM 3h), Pitstop 2 (30 µM 45 min), or diamide (500 µM 45 min), fixed and stained for clathrin (magenta on overlay) and α2-spectrin (green on overlay). Scale bars, 5 µm. E. SIM images of the proximal axon of neurons treated with vehicle (0.1% DMSO 3h), swinholide A (100 nM 3h), Pitstop 2 (30 µM 45 min), or diamide (500 µM 45 min), unroofed and fixed, then stained for clathrin (magenta on overlay) and α2-spectrin (green on overlay). Scale bars, 5 µm. F. Effect of drugs on the density of CCPs along the AIS as quantified from SIM images of intact neurons (n=20, 18, 24, and 18 cells for Ctrl, swinholide A, Pitstop 2, and diamide conditions, respectively). G. Effect of drugs on the density of CCPs along the AIS as quantified from SIM images of unroofed neurons (n=16, 10, 16, and 9 cells for Ctrl, swinholide A, Pitstop 2, and diamide conditions, respectively). H. Effect of drugs on the density of CCPs along the AIS as quantified from PREM views of unroofed neurons (n=25, 8, 11, and 5 images for Ctrl, swinholide A, Pitstop 2, and diamide conditions, respectively). I. PREM views of the proximal axon of unroofed neurons after treatment with swinholide A (100 nM 3h), Pitstop 2 (30 µM 15 min), and diamide (500 µM 15 min), with CCPs colored in magenta on the main image. Zoomed insets correspond to the area highlighted on the main image. Scale bars, 500 nm (main images), 200 nm (insets).

Next, we treated neuronal cultures with the same set of drugs that were used on fibroblasts to target CCP endocytosis (Pitstop 2), actin filaments (swinholide A) and spectrin mesh (diamide). In this case, we first evaluated the density of CCPs at the AIS of treated neurons using SIM of intact and unroofed neurons labeled for clathrin and α2-spectrin (Figure 4, D-E). Pitstop 2 treatment (30 µM for 45 min) resulted in an increase of the CCP density at the AIS compared to controls (from 1.9 ± 0.17 to 2.7 ± 0.17 CCP/µm^2^ for intact neurons, 1.2 ± 0.09 to 2.1 ± 0.14 for unroofed neurons), whereas swinholide A (100 nM for 3h) did not significantly change the density of CCPs (1.8 ± 0.12 and 1.4 ± 0.10 CCP/µm^2^ for intact and unroofed neurons, respectively; Figure 4, F-G). Diamide (500 µM for 45 min) resulted in a visibly patchier appearance of the α2-spectrin mesh at the AIS of unroofed neurons (Figure 4, E) and led to an increased density of CCPs (2.5 ± 0.10 and 2.12 ± 0.21 CCP/µm^2^ for intact and unroofed neurons, respectively; Figure 4, F-G). Quantification of PREM views of treated and unroofed neurons confirmed these results, with Pitstop 2 and diamide A treatment resulting in an increased density of CCPs (from 1.9 ±017 for control to 3.9 ± 0.23, 2.7 ± 0.25, and 4.1 ± 0.44 CCP/µm^2^ for Pitstop 2, swinholide A, and diamide, respectively; Figure 4, H). High-magnification PREM views also resolved the lack of actin filaments in swinholide A-treated neurons and the profound disorganization of the spectrin mesh into connected aggregates caused by diamide (Figure 4, I). Overall, perturbation experiments showed that while CCP formation seems less sensitive to actin disassembly in the AIS compared to fibroblasts. Disorganization of the spectrin mesh increases the density of CCPs at the AIS, suggesting that the disorganization of the periodic spectrin scaffold and its defined clearings makes the plasma membrane more accessible, resulting in the assembly of more CCPs.

### Endocytic cargo concentrates in CCPs at the surface of the AIS

There is no direct correlation between the CCP density at the plasma membrane and the amount of endocytosis occurring in a cell: for example, Pitstop 2 treatment leads to more stalled CCPs and completely inhibits endocytosis (von Kleist et al., 2011), as we verified in Rat2 fibroblasts. Thus, beyond its effect on CCP density, we next sought to directly assess the extent of clathrin-mediated endocytosis after perturbation of the actin-spectrin submembrane scaffold. In fibroblast cells, fluorescent transferrin uptake is a straightforward way to assess clathrin-mediated endocytosis. We first verified that knockdown of α2- and/or ß2-spectrin using siRNAs in Rat2 cells does not downregulate the expression of the transferrin receptor by Western blot (Supporting Figure 3, D). Knockdown of α2- and/or ß2-spectrin, which increase the density of CCPs, also significantly increased the uptake of transferrin from a normalized transferrin fluorescence value of 1.0 ± 0.04 for control to 1.38 ± 0.07, 1.37 ± 0.10, and 1.89 ± 0.09 for knockdown of α2-, ß2-, or α2- and ß2-spectrin, respectively (Figure 5, A-B). By contrast, treatment of fibroblasts with Pitstop 2 and swinholide A, which also increase the density of CCPs, efficiently inhibited the endocytosis of fluorescent transferrin (down to 0.09 ± 0.004 for Pitstop 2 and 0.42 ± 0.04 for swinholide A; Figure 5, C).

We then aimed to assess endocytosis at the AIS of neurons by visualizing the uptake of fluorescent cargo. Incubation of neuronal cultures with fluorescent transferrin for up to 1h resulted in the accumulation of fluorescent puncta and tubules within the cell body and dendrites of neurons, but no endocytosed transferrin could be detected along the proximal axon (Figure 5, D). This is consistent with the literature and the reported absence of transferrin receptors from the axon (Cameron et al., 1991; Farías et al., 2012; Parton et al., 1992). To detect endocytic processes in all neuronal compartments independently of any specific receptor, we thus used fluorescent 10 kDa dextran as a fluid phase marker able to enter cells via clathrin-dependent and independent endocytosis (Li et al., 2015). After 5 to 30 min of incubation at 50 µg/mL, fluorescent dextran clusters were readily detected along the dendrites and the proximal axon (Figure 5, E). Close inspection of SIM images revealed that a fraction of these dextran clusters along the proximal axon of intact neurons were colocalized with CCPs, an observation confirmed by averaged images centered on CCPs, which showed the selective enrichment in dextran at CCPs (Figure 5, F). To verify that a significant proportion of dextran clusters were not found in endosomes, but trapped at the cell surface in CCPs after 30 min of dextran feeding, we unroofed neurons just before fixation to remove the intracellular compartments. SIM images showed that most of the dextran clusters now colocalized with CCPs after unroofing (Figure 5, G). We confirmed that a major population of dextran clusters were still present at CCPs after 30 min of feeding by obtaining correlative SIM-PREM images of unroofed neurons, which showed colocalization of fluorescent dextran clusters at CCPs on PREM views (Figure 5, H). These experiments suggest that despite the presence of a dense population of CCPs in spectrin mesh clearings along the AIS, these CCPs are not efficiently resulting in endocytosis, with endocytic marker clusters being found at the cell surface even after a prolonged incubation.

**Figure 5.**
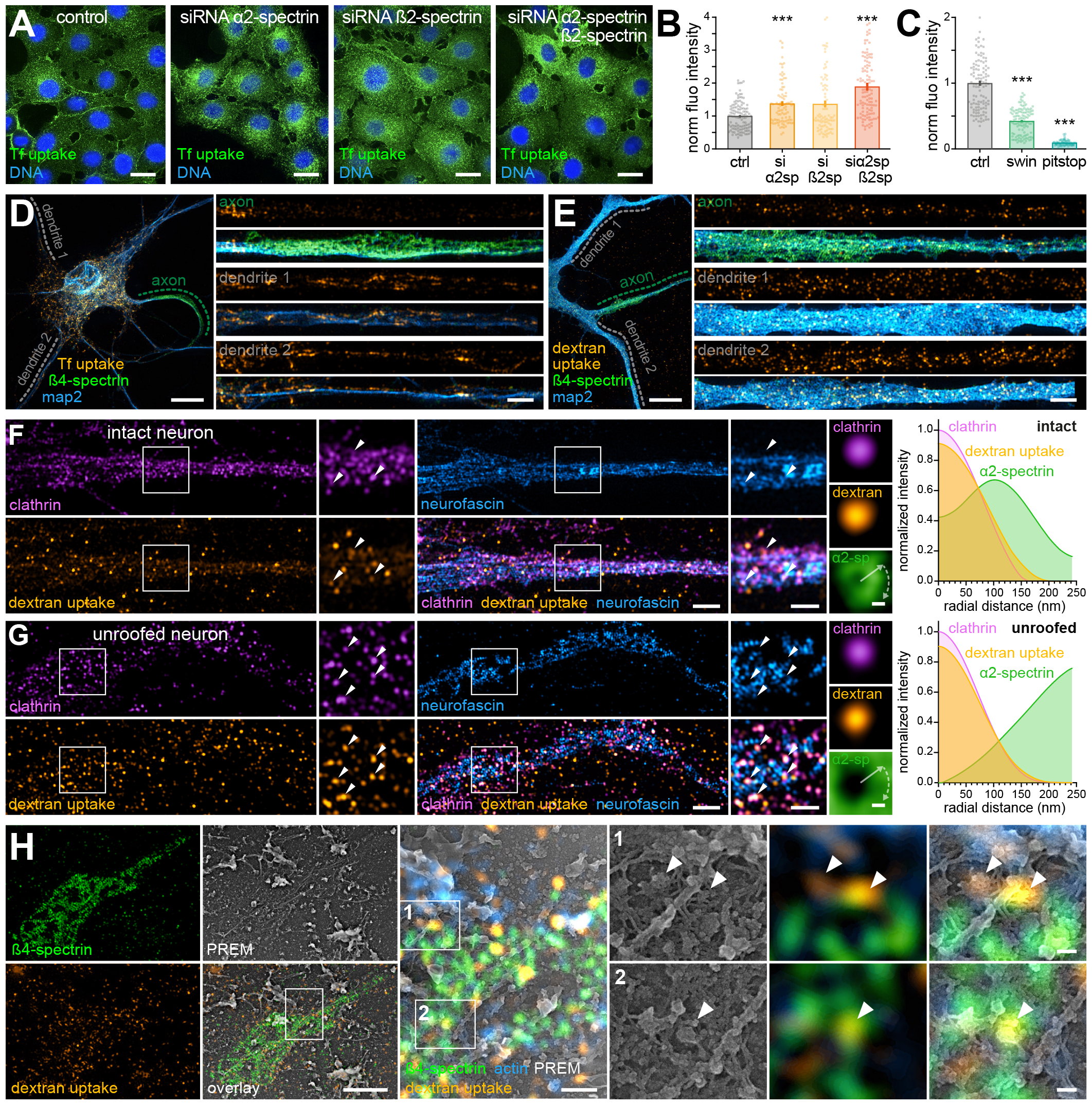
Endocytic cargo concentrates in CCPs at the surface of the AIS. A. Super-resolution spinning disk microscopy images of transferrin-Alexa Fluor 488 uptake (Tf uptake, 20 µg/µL for 10 min, green) in Rat2 fibroblasts transfected with control siRNAs or against α2-spectrin, ß2-spectrin, or α2- and ß2-spectrin (from left to right), fixed and labeled with DAPI (blue). Scale bars, 10 µm. B. Effect of spectrins knockdown on the intensity of Tf uptake in Rat2 fibroblasts, as quantified from confocal images (n=115, 91, 87, and 113 images for Ctrl and siRNA against α2, ß2, and α2/ß2-spectrin, respectively). C. Effect of drugs on the intensity of Tf uptake in Rat2 fibroblasts, as quantified from spinning-disk confocal images (n=115, 116, and 70 images for Ctrl, swinholide A, and pitstop2, respectively). D. SIM image of a neuron fed for 1h with transferrin-Alexa Fluor 647 (orange), fixed, and stained for map2 (dendrites, blue) and ß4-spectrin (AIS, green). Images on the right are straightened zooms along the dendrites and axon highlighted on the main image. Tf uptake is detected along the dendrites, but absent from the AIS and proximal axon. Scale bar, 10 µm (main image), 2 µm (zooms). E. SIM image of a neuron fed for 30 min with dextran-Alexa Fluor 555 (orange), fixed, and stained for map2 (dendrites, blue) and ß4-spectrin (AIS, green). Images on the right are straightened zooms along the dendrites and axon highlighted on the main image. Dextran uptake is detected along the dendrites as well as along the AIS and proximal axon. Scale bar, 10 µm (main image), 2 µm (zooms). F. Left, SIM image of the AIS of a neuron fed with dextran-AF555 (yellow), fixed, and stained for clathrin (magenta) and neurofascin (blue). Some dextran clusters colocalize with CCPs (arrowheads). Right, average images (n=591 pits) centered on CCPs showing the distribution of dextran (orange), clathrin (magenta) and α2-spectrin (green). The graph shows the radial intensity profile corresponding to the average images, with the dextran clusters colocalizing with CCPs in an area devoid of α2-spectrin. Scale bars, 1 µm (AIS images), 100 nm (average images). G. Left, SIM image of the AIS of a neuron fed with dextran-AF555 (yellow), unroofed, fixed, and stained for clathrin (magenta) and neurofascin (blue). Most dextran clusters colocalize with CCPs (arrowheads). Right, average images (n=562 pits) centered on CCPs showing the distribution of dextran (orange), clathrin (magenta) and α2-spectrin (green). The graph shows the radial intensity profile corresponding to the average images (gray arrow), with dextran cluster colocalizing with CCPs in an area devoid of α2-spectrin. Scale bars, 1 µm (AIS images), 100 nm (average images). H. Correlative SIM-PREM image of the AIS of a neuron fed with dextran-AF555 (orange), unroofed, fixed, and stained for ß4-spectrin (green) and actin (blue). Zoomed overlays on the right show dextran clusters found at CCPs in areas devoid of spectrin labeling (arrowheads). Scale bars, 5 µm (isolated SIM channels), 2 µm (main overlay), 100 nm (zooms).

### Live-cell super-resolution microscopy reveals that CCPs are static along the AIS and proximal axon

If CCPs are present at the AIS but rarely result in scission and conclusive endocytosis, they should be long-lived and seen as stable structures by live-cell imaging. To visualize the dynamics of CCPs together with the periodic actin-spectrin scaffold along the submembrane of the proximal axon, we used livecell TIRF-SIM (Kner et al., 2009) of neurons transfected with clathrin light chain (CLC)-mCherry and EGFP-ß2-spectrin (Figure 6, A; Supporting Movie 1). The ∼110 nm lateral resolution of TIRF-SIM could clearly resolve individual CCPs and the 190-nm periodicity of ß2-spectrin along the AIS and proximal axon, and low intensity illumination coupled to deep learning-based denoising allowed to image for tens of frames every 20s. The resulting movie (Supporting Movie 1) and subsequent analysis with kymographs (static CCPs appear as vertical lines, Figure 6, B) or time-color coded projections (static CCPs appear as white dots, Figure 6, C) clearly showed the high stability of CCPs at the AIS compared to the dendrites and cell body of the same neuron, with most AIS CCPs being present across the 20 min of imaging. Temporal autocorrelation analysis of CLC-mCherry intensity profiles from movies of several neurons confirmed this visual impression, with a higher temporal persistence of CLC-mCherry at the AIS compared to the cell body and dendrites (Figure 6, D).

**Figure 6:**
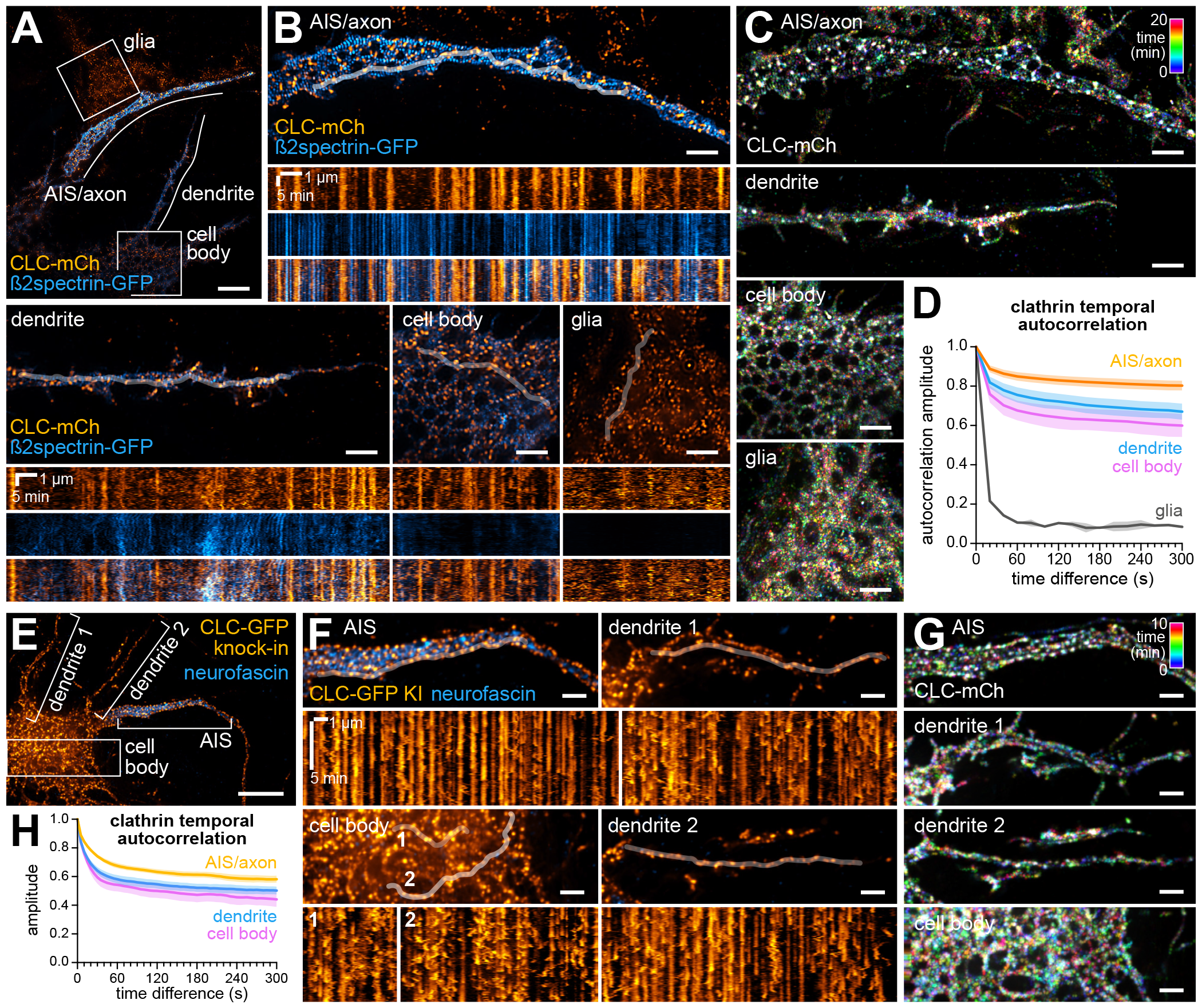
CCPs are static along the AIS and proximal axon. A. Individual frame from Supplementary Movie 1 showing a neuron transfected with ß2spectrin-GFP (blue) and CLC-mCh (orange) and imaged by TIRF-SIM every 20s for 20 min. Scale bar, 5 µm. B. Zooms on areas highlighted in A, with corresponding kymographs (highlighted white lines) along the AIS and proximal axon (top), along a dendrite (bottom left), inside the cell body (bottom center) and a neighboring glial cell (bottom right). Spectrin bands appear as vertical lines spaced by 190 nanometers in blue, while static pits appear as vertical lines and dynamic pits as dots in orange. Scale bars, 2 µm (images), 1 µm (kymograph horizontal), 5 min (kymograph vertical). C. Time color-coded projection of the CLC-mCh channel for the neuronal compartments shown in the panels of B: static pits appear white, while dynamic pits are colored due to their transient appearance. Scale bars, 2 µm. D. Temporal autocorrelation of the clathrin dynamics from tracings within the different compartments of 3 neurons and one glia, showing the relative immobility of CLC in the axon (orange, n=5 tracings) compared to the dendrites (blue, n=5), cell body (magenta, n=4) and within a glial cell (gray, n=2). E. Individual frame from Supplementary Movie 2 showing a neuron with CLC endogenously tagged with GFP (orange), and extracellular staining for neurofascin highlighting the AIS (blue), imaged by SoRa spinning disk every 5s for 10 min. Scale bar, 10 µm. F. Zooms on areas highlighted in E, with corresponding kymographs (highlighted white lines) along the AIS (top left), along two dendrites (top and bottom right), and inside the cell body (bottom left). Static CCPs appear as vertical lines, with putative endocytic events appearing as discontinuities of characteristic J-shaped traces. Scale bars, 2 µm (image), 1 µm (kymograph horizontal), 5 min (kymograph vertical). G. Time color-coded projection of CLC-GFP for the neuronal compartments shown in the panels of B: static pits appear white, while dynamic pits are colored due to their transient appearance. Scale bars, 2 µm. H. Temporal autocorrelation of the clathrin dynamics from tracings within the different compartments of 2 neurons, showing the relative immobility of CLC in the axon (orange, n=5 tracings) compared to the dendrites (blue, n=9) and cell body (magenta, n=7).

To confirm the specific stability of CCPs at the AIS of neurons, and avoid potential stabilizing effects of overexpressed clathrin or spectrin, we next used a modified AAV-mediated HiUGE strategy (Ogawa et al., 2023; Bingham et al., 2023) to endogenously tag clathrin light chain with EGFP in cultured hippocampal neurons (CLC-GFP KI), and imaged clathrin dynamics using super-resolution spinning disk microscopy (Figure 6, E; Supporting Movie 2). The enhanced temporal resolution (one frame every 5s for 10 min) allowed us to discern scission events for both static pits at the AIS (appearing as interruptions in the vertical traces on kymographs) and transient pits in dendrites (appearing as J-shaped traces in kymographs; Figure 6, F). Here again, kymographs and time color-coded projections (Figure 6, G) showed that endogenously-tagged CCPs were more stable at the AIS compared to dendrites and the cell body, an observation confirmed by temporal autocorrelation analysis of CLC-GFP KI from several neurons (Figure 6, H). Overall, these live-cell imaging experiments demonstrate that CCPs are specifically stabilized at the AIS of neurons, consistent with the restrained endocytic activity observed using dextran uptake.

### Endocytosis at the AIS can be triggered by disassembling the spectrin mesh or elevating neuronal activity

At the AIS, CCPs that form in the clearings of the spectrin mesh thus appear to be “frozen”, long-lived structures that do not readily engage in endocytosis. We then wanted to test if perturbation of the actin-spectrin mesh, in addition to increasing the density of CCPs by increasing membrane accessibility, could result in more endocytosis downstream of this increase. We treated neurons with drugs Pitstop 2, swinholide A, and diamide, combining treatments with a dextran uptake assay. Neurons were kept intact before fixation, labeling, and imaging by SIM (Figure 7, A and C) or unroofed before fixation (Figure 7, B and D). Inhibition of CCP scission with Pitstop 2 resulted in a decreased density of dextran clusters, while actin disassembly by swinholide A did not significantly change this density in intact neurons (from 0.66 ± 0.05 clusters/µm^2^ in controls to 0.35 ± 0.04 clusters/µm^2^ for Pitstop 2 and 0.52 ± 0.04 clusters/µm^2^ for swinholide A in intact neurons; Figure 7, C). Both Pitstop 2 and swinholide A did not affect the density of surface dextran clusters after unroofing, suggesting that the decrease in intact neurons for Pitstop 2 was due to inhibited endocytosis (0.72 ± 0.08, 0.70 ± 0.05, 0.64 ± 0.07 for control, Pitstop 2 and swinholide A in unroofed neurons, respectively; Figure 7, D). Interestingly, diamide treatment resulted in an increased density of dextran clusters both in intact and unroofed neurons, indicating enhanced formation of CCPs and increased endocytosis after disorganization of the spectrin mesh at the AIS (0.97 ± 0.07 and 1.19 ± 0.16 clusters/µm^2^ after diamide treatment in intact and unroofed neurons, respectively; Figure 7, C-D). Disorganization of the actin-spectrin mesh thus allows for the formation of more CCPs at the AIS membrane, and results in increased endocytosis.

**Figure 7:**
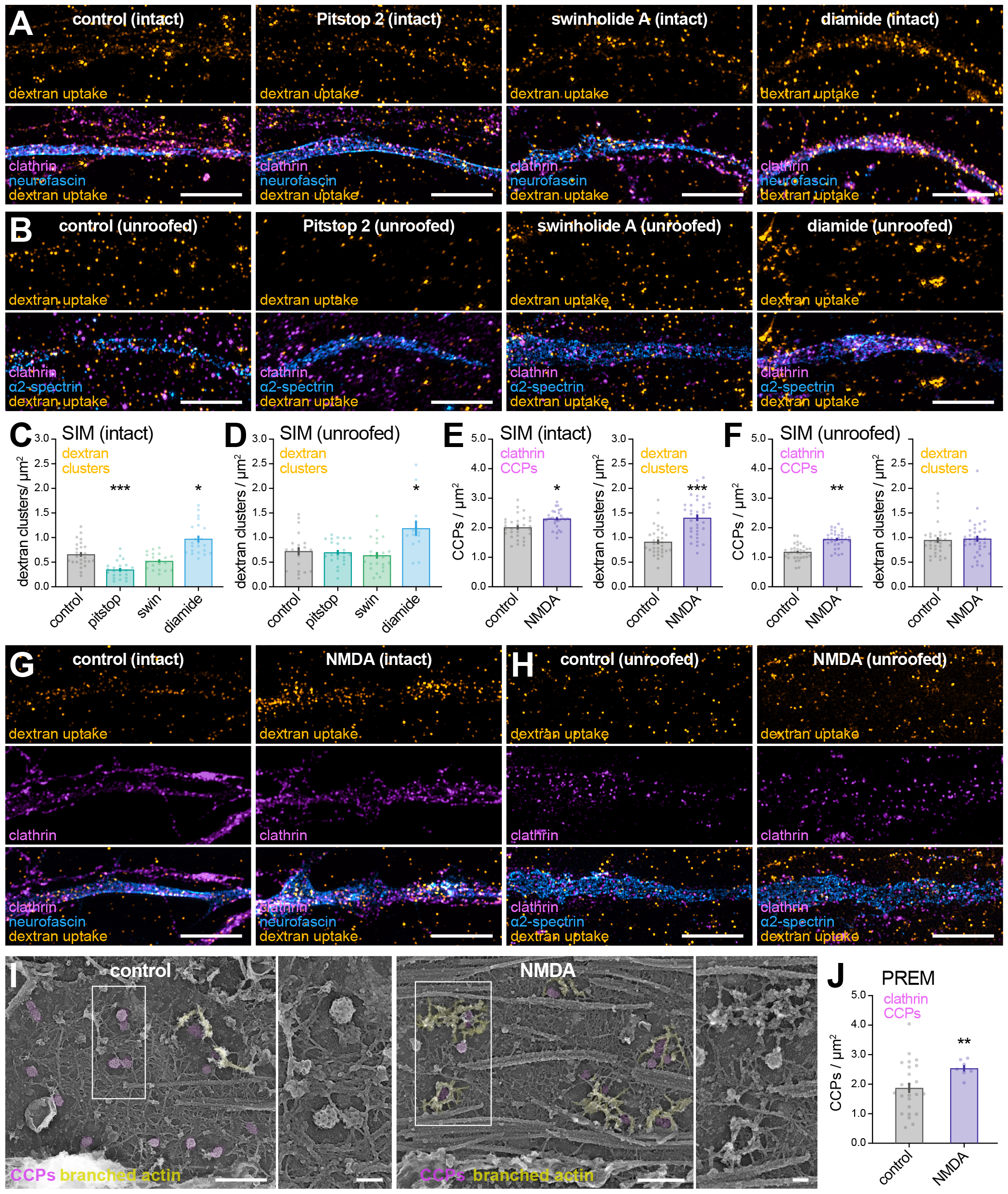
Endocytosis at the AIS can be triggered by disassembling the spectrin mesh or elevating neuronal activity. A. SIM images of the AIS of neurons treated with vehicle (0.1% DMSO 3h), swinholide A (100 nM 3h), Pitstop 2 (30 µM 45 min), or diamide (500 µM 45 min), fed with dextran-AF555 (yellow), fixed, and stained for clathrin (magenta) and neurofascin (blue). Scale bars, 5 µm. B. SIM images of the AIS of neurons treated with vehicle (0.1% DMSO 3h), swinholide A (100 nM 3h), Pitstop 2 (30 µM 45 min), and diamide (500 µM 45 min), fed with dextran-AF555 (yellow), unroofed, fixed, and stained for clathrin (magenta) and neurofascin (blue). Scale bars, 5 µm. C. Effect of drugs on the density of dextran clusters density along the AIS as quantified from SIM images of intact neurons (n=26, 20, 22, and 22 cells for Ctrl, swinholide A, Pitstop 2, and diamide conditions, respectively). D. Effect of drugs on the density of dextran clusters density along the AIS as quantified from SIM images of unroofed neurons (n=21, 23, 23, and 13 cells for Ctrl, swinholide A, Pitstop 2, and diamide conditions, respectively). E. Effect of NMDA-mediated elevation of activity on the CCPs (left graph) and dextran clusters (right graph) density along the AIS as quantified from SIM images of intact neurons (n=31, 31, 32, and 34 cells for Ctrl CCPs, NMDA CCPs, Ctrl dextran, and NMDA dextran conditions, respectively). F. Effect of NMDA-mediated elevation of activity on the CCPs (left graph) and dextran clusters (right graph) density along the AIS as quantified from SIM images of unroofed neurons (n=33, 32, 32, and 35 cells for Ctrl CCPs, NMDA CCPs, Ctrl dextran, and NMDA dextran conditions, respectively). G. SIM images of the AIS of neurons treated with vehicle (0.1% DMSO), or NMDA (50 µm 4 min followed by 30 min rest), fed with dextran-AF555 (yellow), fixed, and stained for clathrin (magenta) and neurofascin (blue). Scale bars, 5 µm. H. SIM images of the AIS of neurons treated with vehicle (0.1% DMSO), or NMDA (50 µm 4 min followed by 30 min rest), fed with dextran-AF555 (yellow), unroofed, fixed, and stained for clathrin (magenta) and neurofascin (blue). Scale bars, 5 µm. I. PREM views of the proximal axon of unroofed neurons after treatment with vehicle or NMDA, with CCPs colored in magenta and branched actin surrounding CCPs colored in yellow on the main image. Zoomed Inset corresponds to the area highlighted on the main image. Scale bars, 1 µm (main images), 200 nm (insets). J. Effect of NMDA-mediated elevation of activity on the density of CCPs along the AIS as quantified from PREM views of unroofed neurons (n=25 and 7 images for Ctrl and NMDA conditions, respectively).

Beyond drug induced perturbations, we finally wondered if a more physiological signal would be able to trigger long-lived AIS CCPs to engage in more efficient endocytosis. It was recently shown that a brief application of NMDA triggered increased neuronal activity and long-term depression (LTD), in turn causing structural AIS plasticity driven by Na_V_1.2 voltage-gated sodium channel endocytosis along the distal AIS (Fréal et al., 2023). We reasoned that this upregulated AIS endocytosis could result from a rearrangement of the actin-spectrin mesh and engagement of CCPs into conclusive endocytosis. Neurons were stimulated with NMDA (50 µM, 4 min) before a 30-minute dextran uptake period in the absence of NMDA. Quantification of CCP and dextran cluster densities from SIM images of intact and unroofed neurons (Figure 7, G-H) showed that NMDA stimulation modestly increased the density of CCPs at the AIS compared to controls (from 2.01 ± 0.07 to 2.30 ± 0.05, from 1.18 ±0.04 to 1.61 ± 0.05 CCP/µm^2^, control to NMDA in intact and unroofed neurons, respectively; Fig. 7, E-F). By contrast, NMDA stimulation induced a significant increase in dextran cluster density in intact neurons (from 0.91 ± 0.05 to 1.40 ± 0.07 clusters/µm^2^ for control and NMDA, respectively; Figure 7, E), whereas it did not change the detected density of surface dextran clusters (from 0.95 ± 0.06 to 0.98 ± 0.06 clusters/µm^2^ for control and NMDA, respectively; Figure 7, F). This suggests that NMDA stimulation can trigger endocytosis from existing CCPs at the AIS membrane. We next visualized the submembrane ultrastructure of NMDA-treated neurons with PREM. High-magnification images of unroofed proximal axons presented a partial disorganization of the spectrin mesh with a more clustered appearance compared to control, and the presence of CCPs with normal morphology (Figure 7, I). We observed an increased presence of branched actin filaments surrounding CCPs within enlarged spectrin clearings, suggesting a role of these filaments in engaging CCPs toward conclusive endocytosis. Quantification of CCP density showed a modest increase in CCPs density along the AIS, consistent with the SIM data (from 1.87 ± 0.17 CCP/µm^2^ in controls to 2.54 ± 0.11 CCP/ µm^2^ after NMDA stimulation; Figure 7, J). Elevated activity is thus able to stimulate endocytosis at the AIS, by unlocking the engagement of long-lived CCPs.

## Discussion

The AIS and proximal axon are neuronal compartments where endocytosis has long been overlooked. Our experiments combining super-resolution optical and electron microscopy with genetic and pharmacological perturbations provide compelling evidence that the AIS contains a population of CCPs with unique properties, allowing endocytosis to be tightly controlled by the AIS submembrane scaffold.

We discovered that in the AIS, CCPs form on bare areas of the plasma membrane that are precisely delimited by a circular clearing of the actin-spectrin undercoat. These ∼300 nm clearings are scattered across the inner surface of the AIS and proximal axon, inserted within the previously described periodic scaffold of actin rings connected by spectrin tetramers (Vassilopoulos et al., 2019). They exhibit a specific organization: actin filaments are present at the border of the clearings, crosslinked by spectrins and ankyrin that form an outer polygonal arrangement. In addition, a single actin filament is often seen to enter the clearing and contact the central CCP. We propose that clearings are structures that allow the formation of CCPs by providing access to the AIS plasma membrane. The necessity for CCPs to form in membrane areas devoid of spectrins was previously shown by diffraction-limited fluorescent microscopy in spectrin-dense regions of fibroblasts (Ghisleni et al., 2020), and along the lateral membrane of epithelial cells (Jenkins et al., 2015) – it makes all the more sense along axons where the spectrin mesh is particularly dense.

This organization of CCPs in clearings defined by the submembrane spectrin scaffold immediately suggests that this scaffold can control the formation of CCPs by defining the location and density of these clearings. This inhibitory role of submembrane spectrins on CCP formation and subsequent endocytosis was proposed for fibroblasts (Ghisleni et al., 2020) and it was shown that endocytosis was negatively controlled by the ankyrin/spectrin domains at the lateral membrane of epithelial cells (Jenkins et al., 2015). Here, we demonstrate that spectrin knockdown or disorganization directly upregulates CCP formation and endocytosis in fibroblasts and along the AIS in neurons, establishing the spectrin mesh as an insulating layer restricting endocytic processes. Beyond the AIS, the periodic actin-spectrin scaffold is likely to regulate endocytosis along the whole axon, as shown by the enhanced endocytosis of cannabinoid receptors after activation in neurons depleted for ß2-spectrin, part of the distal axon periodic scaffold (Zhou et al., 2019).

Our results suggest that another layer of regulation is present besides the insulating effect of the spectrin mesh. We found that CCPs that do form in mesh clearings along the AIS are unusually long-lived and do not readily engage in conclusive endocytosis, accumulating cargoes at the AIS surface. Actin has been shown to be important for the late stages of clathrin-mediated endocytosis (Loebrich, 2014), as confirmed by the presence of numerous stalled CCPs in fibroblasts after swinholide A treatment. What could explain the “frozen” property of CCPs at the AIS? We speculate that the dense spectrin mesh could prevent the polymerization of actin around pits, impeding the progression of CCPs toward scission and conclusive endocytosis.

These two layers of regulation allow us to explain the observed effects of different perturbations on CCP formation and endocytosis at the AIS: swinholide A disassemble actin structures, but cannot fully disassemble the periodic spectrin scaffold at the AIS (Vassilopoulos et al., 2019). In addition, it does not induce the accumulation of low-curvature clathrin-coated structures by blocking the late stage of endocytosis like in fibroblasts, as this block is already occurring at the AIS in control conditions and neurons do not form flat clathrin lattices (Moulay et al., 2020). Depletion of the spectrin mesh by knockdown and its disorganization by diamide increase the accessible membrane area and increase the number of CCPs, while releasing actin from the submembrane mesh that can help CCPs to engage in conclusive endocytosis, resulting in more internalized cargo. Finally, NMDAR activation is a more subtle and physiological intervention that does not profoundly alter the spectrin mesh, but could allow the formation of branched actin interacting with CCPs, resulting in the engagement of existing CCPs into conclusive endocytosis (Engqvist-Goldstein et al., 2004; Vassilopoulos et al., 2014).

What could be the functional relevance of this tight regulation of endocytosis at the AIS by the submembrane spectrin scaffold? Firstly, static CCPs could help sequester membrane proteins that do not bind to AIS-specific scaffolds like ankyrin G, priming them for a low but continuous process of endocytic retrieval and degradation as a sorting mechanism (Eichel et al., 2022). Secondly, the presence of pre-arranged, static endocytic spots in the dense spectrin mesh likely facilitates the rearrangement of AIS membrane proteins that are bound to the remarkably stable ankyrin/spectrin scaffold (Brachet et al., 2010): endocytosis of sodium channels or neurofascin during morphological plasticity (Fréal et al., 2023) would only require detachment from ankyrin G – a process likely to be phosphorylation-dependent (Tuvia et al., 1997; Bréchet et al., 2008) – and diffusion to the nearest CCP. An “unfreezing” mechanism such as the one we evidenced after NMDA treatment would thus trigger efficient endocytosis from these CCPs, allowing to modify the AIS length or position (Yamada and Kuba, 2016).

Overall, our results support a model where CCPs form in newly-described structures, “clearings” of the actin-spectrin mesh along the AIS, and are then stabilized as long-lived structures with moderate endocytic activity at steady-state. Efficient endocytosis can be triggered by physiological signals, allowing to rearrange the AIS membrane content or internalize extra-cellular components. We look forward to further explore the mechanisms and roles of this regulated process, which is likely to have important physiological and pathological implications for neuronal organization and function.

## Acknowledgments

We would like to acknowledge funding by the Agence Nationale pour la Recherche (ANR-20-CE16-021-03 to C.L. and S.V., ANR-21-CE13-0018-01 to SV), Fédération pour la Recherche sur le Cerveau (AOE 16 “Espoir en tête” 2021) to C.L., Sorbonne Université, INSERM, Association Institut de Myologie core funding to S.V. We would like to thank the Neuro-Cellular Imaging Service and Nikon Center for Neuro-NanoImaging at INP, with funding from CPER-FEDER (PlateForme NeuroTimone PA0014842), the Institut Marseille Imaging and NeuroMarseille for complementary equipment funding from Excellence Initiative of Aix-Marseille University – A*MIDEX, a French “Investissements d’Avenir” program (AMX-19-IET-002), and NeuroSchool for end-of-PhD funding. We would also like to thank the IBPS electron microscopy platform (Sorbonne University, Paris, France) and the MyoVector facility from the Institute of Myology (Paris, France). We would like to thank Subhojit Roy and Marc Bitoun for insightful discussions, as well as Damaris Lorenzo for the gift of the EGFP-ß2-spectrin plasmid.

## Author Contributions

Conceptualization: C.L., S.V.; Methodology: F.W., S.M., J.L., N.J., C.L., S.V.; Formal analysis: F.W., S.M., C.L., S.V.; Investigation: F.W., S.M., J.L., G.M., F.B.-R., F.P., S.B.-Z., M.-J.P., C.L., S.V.; Resources: N.J.; Writing-Original Draft: S.M., C.L., S.V.; Writing-Review and Editing: F.W., S.M., J.L., M.-J.P., C.L., S.V.; Visualization: F.W., S.M., C.L, S.V.; Supervision, Project Administration: C.L., S.V.; Funding acquisition: C.L., S.V.

## Methods

### Animals and neuronal cultures

The use of Wistar rats followed the guidelines established by the European Animal Care and Use Committee (86/609/CEE) and was approved by the local ethics committee (agreement D13-055-8). Rat hippocampal neurons were cultured following the Banker method, above a feeder glia layer (Kaech and Banker, 2006). Rapidly, 12 or 18 mm-diameter round #1.5H coverslips were affixed with paraffin dots as spacers, then treated with poly-L-lysine (Sigma-Aldrich #P2636). Hippocampi from E18 rat pups were dissected and homogenized by trypsin treatment followed by mechanical trituration and seeded on the coverslips at a density of 4,000-8,000 cells/cm^2^ for 3h in serum-containing plating medium. Coverslips were then transferred, cells down, to petri dishes containing confluent glia cultures conditioned in B27-supplemented (Thermo Fisher Scientific #17504044) neurobasal medium (NB, Thermo Fisher Scientific #21103049), and cultured in these dishes for up to 4 weeks. For this work, neurons were fixed at 13-16 days in vitro (div), a stage where they exhibit a periodic actin/spectrin scaffold along virtually all axons (Xu et al., 2013).

### Fibroblast cultures and siRNA-mediated knockdown of spectrins

Rat2 fibroblasts were seeded on 18-mm glass coverslips to 80 % confluency in DMEM containing 10 % FCS, and they were incubated overnight at 37°C. Prior to transfection, cells were washed in PBS. Complexes of Lipofectamine 3000 (Thermo Fisher Scientific) and siRNA (corresponding to the shRNA sequences detailed below) were then introduced to the cells following the manufacture protocol (Thermo Fisher Scientific). The cells were left to incubate for 48 hours.

The validation of siRNAs knock-down in fibroblasts was performed by Western blot. Rat2 cultures were lysed in a buffer (50 mM Tris-HCl, pH 7.5, 0.15 M NaCl, 1 mM EDTA, 1% NP-40) supplemented with a protein inhibitor cocktail 1:100 (Sigma-Aldrich). Samples denatured by 3 min boiling in Laemmli buffer were separated by electrophoresis on 4-12% bis-acrylamide gel (Thermo Fisher Scientific), then transferred to 0.45 µm nitrocellulose membranes (Thermo Fisher Scientific) and labeled with a primary antibody then a secondary antibody coupled to horseradish peroxidase (Trueblot immunoglobulin G HRP, Rockland). Proteins in samples were detected using Immobilon Western Chemiluminescent HRP Substrate (Sigma-Aldrich) and image acquisition was performed on a G-Box (Ozyme). The following primary antibodies were used to validate siRNA knock-down in fibroblasts: anti-α2-spectrin (mouse IgG2b clone D8B7, Biolegend #803201, 1:1000), anti-β2-spectrin (BD Biosciences #612563, mouse IgG1 against residues 2101–2189 of human β2-spectrin, 1:100), anti-clathrin (rabbit polyclonal anti-CHC, Abcam #ab21679, 1:1000), anti-TfR (mouse IgG1, Invitrogen #13-6800, 1:1000); anti-GADPH (mouse IgG1, Santa Cruz #sc-47724, 1:1000) was used as a loading control.

### Plasmid constructs and neuronal culture transfection

Transfections were performed using the following constructs: clathrin-GFP and clathrin-mCherry plasmids were a gift from Subhojit Roy (Ganguly et al., 2021). The EGFP-β2-spectrin plasmid was a gift from Damaris Lorenzo (Cousin et al., 2021). The spectrin-silencing shRNA plasmids were constructed in the pRFP-C-RS backbone (Origene). These plasmids allow cloning and expression of shRNA under a U6 promoter and co-expression of the TurboRFP red fluorescent protein under the control of a CMV promoter. Insertion of the shRNA sequences was performed using the SLIC technique using overlapping single strand oligonucleotides (Jeong et al., 2012).

The sequences for each shRNA were:

α2-spectrin: **g**ATCAGTTTGTGGAAGAAACTTtcaa-gagAAGTTTCTTCCACAAACTGAT

β2-spectrin: **g**CAGAAGAGATCTCCAACTACAtcaagagTG-TAGTTGGAGATCTCTTCTG

β4-spectrin: **g**CACTGGATAGCCGAGAAGGtcaagag-CCTTCTCGGCTATCCAGTG

Luc (negative control): **g**CGCTGAGTACTTCGAAAT-GTCtcaagagGACATTTCGAAGTACTCAGCG

(First g in bold-lowercase: additional g for efficient U6 transcription, underscore-lowercase: shRNA loop. A shRNA directed against the luciferase coding sequence was used as negative control).

Plasmid constructs (0.25 to 0.5 µg) per 18-mm coverslip were incubated with 1 µL of Lipofectamine 3000 (Thermo Fisher Scientific #L3000001) in 50 µL NB for 20 min at room temperature for liposomes to form, before being added to 10 div cultured hippocampal neurons, coverslips flipped cells up in FB12 dishes containing preheated NB. Neurons were incubated for 30 min at 37°C with 5% CO_2_. After incubation, the neurons were returned to their origin culture dishes and left incubating for 2.5 to 3 days before imaging.

### AAV production and titration

AAV viral vectors were prepared by triple transfection in HEK-293 cells using polyethyleneimine transfection agent: pSMD2 AAV vector plasmid containing the shRNA sequences detailed above, pXX6 plasmid coding for the viral sequences essential for AAV production, and p0009 plasmid coding for serotype 9 capsid. Vector particles were purified on an iodixanol gradient and concentrated on Amicon Ultra-15 100K columns (Merck-Millipore). The AAV vectors were titrated as vg per milliliter by quantitative real-time PCR using ITR2 (inverted terminal repeats) specific primers at a 60°C annealing temperature (forward 5′-CTCCATCACTAGGGGTTCCTTG-3′ and reverse 5′-GTAGATAAGTAGCATGGC-3′) and the MGB Taqman probe 5′-TAGTTAATGATTAACCC-3.

### AAV infection for shRNA-mediated knockdown in neuronal cultures

Spectrin-silencing shRNAs in adeno-associated virus vectors (AAV) were produced and titrated by the AAV production service of Institut de Myologie (Paris). Vectors identifiers AAV9-U6SCR (p05.1 Origen scrambled sequence GAGAGCCAGATTCAATCTGACGACTATGG), AAV9-U6beta4 / p10.1 (Hedstrom et al., 2007), AAV9-U6alpha2 / p09.1 (Galiano et al., 2012), AAV9-U6-shRNA-α2+ß4-spectrin / p11.1 (both combined) targeted respectively no protein (scrambled control), β4-spectrin, α2-spectrin, α2- and β4-spectrin. The validation of shRNAs knock-down in neurons was performed by Western blot. Cells infected with an AAV-shRNAs were resuspended in a loading buffer with DTT, diluted to 50% in ultrapure water. The samples were then heated to 90°C for 3 min. After extensive washes, samples were resolved by 7.5% sodium dodecyl sulfate–polyacrylamide gel electrophoresis, and immunoprobed with one of anti-α2-spectrin (Biolegend #803201, mouse IgG2b clone D8B7, 1:1000), anti-β2-spectrin (BD Biosciences #612563, mouse IgG1 against residues 2101–2189 of human β2-spectrin, 1:100), anti-β4-spectrin Cter (from Matthew Rasband, Baylor College of Medicine Austin TX, rabbit against residues 2237-2236, 1:100), an anti-α tubulin (Sigma-Aldrich #T5168, mouse IgG1 clone B-5-1-2, 1:1500) was used as control. The shRNAs effect on the periodicity of the axonal actin-spectrin scaffold was evaluated by fluorescence intensity quantification and autocorrelation of intensity profiles along axonal segments (https://github.com/cleterrier/Process_Profiles).

### Endogenous clathrin tagging (knock-in)

We used a rat optimized version of the HiUGE method (Gao et al., 2019; Ogawa et al., 2023), with an adapted Donor Recognition Sequence (DRS) GCGATCGTAATCACCCGAGT-GGG. We used the homologous sequence of the protospacer used by Gao et al. to cut the clathrin light chain A subunit gene (Clta).as close as possible to the stop codon (5 nucleotides before):

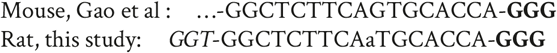

In italics: Additional nucleotides in 5’ to have a 20-nucleotide length protospacer sequence. In bold the PAM sequence on the Clta gene. In lowercase the nucleotide that differs from the mouse sequence.

AAVs were produced by the standard triple transfection in HEK293T in the AAV9/PHP.S serotype. Small scale AAV preparations were obtained by mild lysis of HEK cells 3 days post transfection in a citrate buffer (38.1 mM citric acid, 74.8 mM NaHCO_3_, 75 mM NaCl, 100 mM MgCl, pH 5) and neutralized subsequently with Tris-HCl pH 9.5. AAV particles were titrated by qPCR with an improved version of the Addgene SYBR green method. In brief, we used a linearized AAV plasmid as standard instead of a supercoiled plasmid and we incorporated a proteinase K step after the DNAse treatment. We usually obtained titers between 10^11^ and 10^12^ part/mL. Hippocampal neurons were triply infected after plating at 0 div and at a density of 12,000 cells/cm^2^ with 10^9^ AAV particles for each 3 different AAVs: Mecp2-Cas9 AAV, clathrin-Cter sgRNA AAV and DRSR2-EGFP-ORF1 Cter HiUGE payload.

### Pharmacological treatments

Treatments were applied on neurons and fibroblasts in their original culture medium at 37°C, 5% CO_2_. Dimethyl sulfoxide (DMSO) 0.1% (from pure DMSO, Sigma-Aldrich #D2650) was used as a control vehicle, Pitstop 2 30 µM (from 30 mM stock in DMSO, Abcam #120687) was applied for 15 min (45 min in neuron feeding experiments), swinholide A 100 nM (from 100 μM stock in DMSO, Sigma-Aldrich #S9810) was applied for 3h, diamide 500 µM (from 1M stock in ultrapure water, Sigma-Aldrich #D3648) was applied for 15 min (45 min in neuron feeding experiments). All stock solutions were stored at −20°C. For NMDA stimulation, 14 div neurons were first incubated 15 min in 500 µL of uncomplemented neurobasal medium (NB) in a 12 wells plate at 37°C 5% CO_2_, then NMDA (from 50 mM stock in ultrapure water, Sigma-Aldrich #M3262), diluted to 100 µM in preheated NB, was added to the wells for a final concentration of 50 µM NMDA. For control experiments, NB without NMDA was used. After 4 min, neurons were gently washed 3 times with 500 µL NB, then returned to their original complemented medium and incubated for 30 min. Neurons were then either fixed and stained as described in the immunocytochemistry section or incubated with an endocytosis probe before as described in the uptake assays section.

### Unroofing and fixation of fibroblasts and neuronal cultures

Unroofing was performed by sonication as previously described (Heuser, 2000). Coverslips were quickly rinsed three times in Ringer+Ca (155 mM NaCl, 3 mM KCl, 3 mM NaH_2_PO_4_, 4.5 mM HEPES, 10 mM glucose, 2 mM CaCl_2_, 1 mM MgCl_2_, pH 7.2), then immersed 10s in Ringer-Ca (155 mM NaCl, 3 mM KCl, 3 mM NaH_2_PO4, 4.5 mM HEPES, 10 mM glucose, 3 mM EGTA, 5 mM MgCl_2_, pH 7.2) containing 0.5 mg/mL poly-L-lysine, then quickly rinsed in Ringer-Ca then unroofed by scanning the coverslip with rapid (2-5s) sonicator pulses at the lowest deliverable power in KHMgE buffer (70 mM KCl, 30 mM HEPES, 5 mM MgCl_2_, 3 mM EGTA, pH 7.2). Unroofed cells were immediately fixed in 4% PFA in KHMgE for 10 min for epifluorescence microscopy, SIM and SMLM, for 45 min for PREM of immunogold-labeled samples, and for 15 min for PREM, correlative SIM-PREM and SMLM-PREM. After fixation and rinses, all neuronal samples for PREM were placed in individual sealed transport wells containing 5 mL of KHMgE with 2% glutaraldehyde and shipped immediately the following day. Samples for correlative light microscopy-PREM were immunolabeled and imaged immediately and kept in KHMgE with 2% sodium azide (Sigma-Aldrich #08591) at 4°C for a maximum of 1.5 days before being shipped in the same manner.

### Fluorescence immunocytochemistry

Fluorescence immunocytochemistry of neuronal cultures for widefield microscopy, SIM and SMLM was performed as in published protocols (Jimenez et al., 2020). Cells were fixed using 4% PFA in PEM buffer (80 mM PIPES pH 6.8, 5 mM EGTA, 2 mM MgCl_2_) for 10 min at room temperature (RT). After rinses in 0.1 M phosphate buffer (PB), neurons were blocked for 2-3h at RT in immunocytochemistry buffer (ICC: 0.22% gelatin, 0.1% Triton X-100 in PB), and incubated with primary antibodies diluted in ICC overnight at 4°C. After rinses in ICC, neurons were incubated with secondary antibodies diluted in ICC for 1h at RT and rinsed. Actin staining was then performed by incubating in fluorescent phalloidin at 0.5 μM for either 1h at RT or overnight at 4°C. Stained coverslips were kept in PB + 0.02% sodium azide at 4°C until imaging by SMLM. For widefield microscopy and SIM, coverslips were mounted in ProLong Glass (Thermo Fisher Scientific #P36980).

Immunofluorescence of unroofed neurons for correlative SIM-PREM was performed using a fast protocol, beginning with 3 rinses in KHMgE buffer, then 30 min blocking in detergent-free buffer (DFB: KHMgE, 1% BSA), 30 min primary antibodies incubation at RT, 3 DFB rinses with 5 min incubation between them, 30 min secondary antibodies incubation at RT, 3 DFB rinses as before, then a KHMgE rinse, keeping the unused coverslips in KHMgE at 4°C until acquired in SIM.

Immunogold labeling of unroofed samples was performed in the same detergent-free solution: samples were blocked for 30 min, incubated for 90 min with the primary antibodies diluted to 1:20, rinsed, incubated two times 20 min with the gold-coupled secondary antibodies, then rinsed.

### Antibodies labeling, probes and other reagents

Rabbit polyclonal anti β4-spectrin antibody (against residues 2237–2256 of human β4-spectrin, 1:500 dilution for immuno-fluorescence IF, 1:20 for immunogold IG) was a gift from Matthew Rasband (Baylor College of Medicine, Austin, TX). Mouse monoclonal anti β2-spectrin (against residues 2101–2189 of human β2-spectrin, 1:100 for IF) was from BD Biosciences (#612563). Mouse monoclonal anti α2s-spectrin (clone D8B7, 1 :100 for IF) was from Biolegend (#803201). Rabbit polyclonal anti 480-kDa ankyrin G (residues 2735–2935 of rat 480-kDa ankyrin G, 1:300 for IF, 1:100 for IG) was a gift from Vann Bennett (Duke University, Durham, NC). Rabbit anti phospho-myosin light chain 2 Thr18/Ser19 (pMLC, 1:50 for IF, 1:20 for IG) was from Cell Signaling Technologies (#3674). Chicken anti-map2 (against residues 2-314 of human map2, 1:1000 for IF) was from Synaptic Systems (#188006). Chicken anti pan-neurofascin (against residues 25-1031 of rat neurofascin, 1:400 for IF) was from R&D System (#AF3235). Rabbit polyclonal anti-clathrin heavy chain (against residues 1650 to C-terminus of human clathrin heavy chain, IF 1:150) was from Abcam (#ab21679). Mouse monoclonal anti-clathrin light chain (clone X22, 1:100 for IF) was from Thermo Fisher Scientific (#MA1-065); we also used clone CON.1 (1:1000 for IF) from Sigma-Aldrich (#c1985). Mouse monoclonal anti-AP2 (1:100 for IF) was from Abcam (#ab2730).

### Secondary antibodies and reagents

Donkey and goat anti-rabbit, anti-mouse, and anti-chicken secondary antibodies conjugated to Alexa Fluor 488, 555, and 647 were from Life Technologies or Jackson ImmunoResearch (1:200-1:400 for IF). Goat anti-rabbit and anti-mouse secondary antibodies conjugated to DNA-PAINT handles were obtained from Massive Photonics (MASSIVE-sdAB 2-PLEX). Goat anti-rabbit and anti-mouse secondary antibodies conjugated to gold nanobeads were from Aurion (15 nm gold, #815011 and #815022, respectively, 1:20 for IG) or Jackson ImmunoResearch (18 nm gold, #111-215-144 and #115-215-146, respectively, 1:15 for IG). For light microscopy, we used Alexa Fluor 488 and Alexa Fluor 647 Plus-conjugated phalloidin from Thermo Fisher Scientific (#A12379 and #A2287, respectively) and Atto488-conjugated phalloidin (#AD48881) from Atto-Tec. For immunogold labeling of actin, unroofed neurons were fixed with 2% PFA in KHMgE, then quenched for 10 min in KHMgE, 100 mM glycine, and 100 mM NH4Cl. After blocking in KHMgE, 1% BSA, they were incubated with phalloidin-Alexa Fluor 488 (0.5 μM) for 45 min, then immunolabeled using an anti-Alexa Fluor 488 primary antibody and a gold coupled goat anti-rabbit secondary antibody as described above for immunogold labeling. Paraformaldehyde (PFA, #15714, stock 32% in water) was from Electron Microscopy Sciences, and glutaraldehyde (#G5882, stock 25% in water) was from Sigma-Aldrich.

### Transferrin and dextran uptake assays

For pulse-chase uptake assay of transferrin internalization in rat fibroblasts, cells were cultured on 12-mm glass coverslips in DMEM and 10% FCS to 80% confluency. Transferrin from human serum conjugated to Alexa Fluor 488 (Thermo Fisher Scientific #T3342) was used at 20 µg/mL for 10 min at 37°C. Cells were acid-stripped (0.2 M Na_2_HPO_4_, 0.1 M citric acid), rinsed three times in PBS, and then fixed in 4 % paraformaldehyde (PFA) for 20 min. Glass slides were mounted with coverslips using Vectashield containing DAPI (Vector Laboratories #H-1200). Regions of interest were defined based on the cell shape. Individual cells were manually outlined and corrected total cell fluorescence = integrated density - (area of selected cell × mean fluorescence of background readings) was calculated using Fiji imaging software.

For transferrin uptake experiments in neurons, cells that were incubated for 1h in uncomplemented Neurobasal medium, then Alexa-Fluor-647-conjugated transferrin was used (Thermo Fisher Scientific, #T23366) diluted at 1:50 in the same medium and incubated for 1h at 37°C and 5% CO_2_. After incubation, the cells were quickly rinsed 3 times with pre-heated medium and fixed with PFA 4% in PEM buffer. For dextran uptake experiments, the same procedure was used, with Alexa Fluor 555-conjugated dextran 10 kDa (Thermo Fisher Scientific #D34679) diluted at 50 µg/mL and incubated for 30 min at 37°C and 5% CO_2_ before rinses and fixation.

### Live-cell imaging probes and AIS staining

All live-cell experiments were performed in live-cell imaging medium (Hibernate E low fluorescence, Brainbits/Thermo Fisher Scientific #NC0285514) supplemented with 3% glucose, 2% B27, and 0.5mM L-glutamine). Live-cell AIS staining was performed using a mouse anti-neurofascin antibody targeting an extracellular epitope (NeuroMab clone A12/18, #75-172) conjugated with CF647 (Mix-N-Stain CF647 antibody labeling kit, Sigma-Aldrich #MX647S50) following the manufacturer protocol. The conjugate was diluted to 1:50 and incubated for 10 min at 37°C and 5% CO_2_ before proceeding to SIM.

### Structured illumination microscopy

SIM of neurons was performed on an N-SIM-S microscope (Nikon Instruments). The N-SIM system uses an inverted Nikon Eclipse Ti2-E microscope with a perfect focus system 4, an integrated Nikon laser launch with 405, 488, 561, and 640 nm solid-state excitation lasers, a 100X NA 1.49 oil objective (SR HP Apo TIRF), a Mad City Labs Nanodrive piezo stage and a Hamamatsu Fusion BT CMOS camera. On fixed cells, after locating a neuron of interest using low-intensity illumination, an image was acquired in 2D-SIM/TIRF-SIM (9 images per image) or 3D-SIM (15 images per z plane) mode. Reconstruction was performed using the N-SIM module in the NIS-Elements software (AR 5.30.05), resulting in a ∼120 nm lateral (all modalities) and ∼250 nm axial resolution (for 3D-SIM).

For live-cell SIM, live-cell imaging medium was used, and the stage was kept at 37°C by a Tokay Hit STX heater. TIRF-SIM movies were captured at a frequency of 0.05 Hz for 15 to 20 min, alternating the 488- and 561-nm illumination to image ß2-spectrin-GFP and CLC-mCherry, respectively. Movies were processed using a deep-learning based denoising module (Enhance.ai in NIS-Elements software, Nikon) using a model trained from 20 pairs of fixed samples imaged at low (10%-20%) and high (100%) laser intensities. Neurons used for training were transfected with CLC-mCherry and EGFP-β2-spectrin, then fixed and mounted in live-cell imaging medium.

For super-resolved spinning disk microscopy of Rat2 fibroblasts, images were acquired using a Nikon Ti2 microscope, driven by Metamorph (Molecular Devices), equipped with a motorized stage and a Yokogawa CSU-W1 spinning disk head coupled with a Prime 95 sCMOS camera (Photometrics) equipped with a 100X oil-immersion objective lens. Super-resolution images were obtained using the LiveSR module (Gataca Systems). DAPI, Alexa Fluor 488, Alexa Fluor 568 and Alexa Fluor 647 were sequentially excited. Z-series from the top to the bottom of fibers were sequentially collected for each channel with a step of 0.1-0.3 µm between each plane.

Live-cell super-resolved spinning-disk imaging of endogenously-tagged clathrin-EGFP in neurons was performed on a Yokogawa CSU-W1 spinning disk microscope equipped with 405, 488, 561, and 640 nm solid-state excitation lasers, a SoRa pixel reassignment module, a Ti2-E stand (Nikon), a 60X NA 1.49 oil objective, and a Fusion BT sCMOS camera (Hamamatsu). Livecell imaging medium was used, and the stage was kept at 37°C by a stage enclosure (Okolab). After capturing a single image of the neurofascin live-cell staining using 647-nm laser excitation, super-resolved images of clathrin-GFP were obtained using 488-nm laser excitation, with the SoRa module 4X magnification lens and a camera binning of 2×2 at a frequency of 0.2 Hz. Movies were processed using a pre-trained deep-learning based denoising module embedded in the NIS-Elements software (Denoise.ai, Nikon).

### SMLM: STORM and PAINT

Both STORM and DNA-PAINT acquisitions were performed on an N-STORM microscope (Nikon Instruments). The N-STORM system uses an Agilent MLC-400B laser launch with 405 nm (50 mW maximum fiber output power), 488 nm (80 mW), 561 mW (80 mW), and 647 nm (125 mW) solid-state lasers, a 100X NA 1.49 objective, and an Ixon DU-897 camera (Andor). After locating an axon using low-intensity epifluorescence illumination, followed by either a STORM or DNA-PAINT acquisition using laser illumination in HiLo (grazing angle) configuration. An astigmatic lens was added to the light path to achieve 3D imaging. For STORM, stained coverslips were mounted in a silicone chamber filled with STORM buffer (Smart Buffer Kit; Abbelight), and 30,000-60,000 images (256 × 256 pixels, 15 ms exposure time) were acquired at 100% 647-nm laser power. Reactivation of fluorophores was performed during acquisition by increasing illumination with the 405-nm laser. For DNA-PAINT, stained coverslips were mounted in a Ludin chamber filled with imaging buffer (Massive Photonics). Imaging strands conjugated to Atto565 and Atto655 (Massive Photonics) were added from a starting concentration of 0.1 nM each, then adjusted to optimize blinking density. 30,000-45,000 images (256×256, 40 ms exposure time) were acquired at 25-50% laser power for 561 nm, 60-100% laser power for 647 nm by alternating frames with 561-nm and 647-nm laser excitation.

For STORM and DNA-PAINT images, acquired stacks were processed using DECODE (Speiser et al., 2021). Briefly, PSFs were modeled using spline fitting in SMAP (Li et al., 2018) and used to simulate sequences of blinking events using characteristics (photon number range and lifetime distribution) inferred from real acquisition data from the N-STORM microscope. A PyTorch model was trained to infer the 3D coordinates and uncertainty of the simulated blinking events and then applied to the experimental acquired sequence (Speiser et al., 2021). The resulting localizations (fitted blinking events) were filtered based on uncertainty, and drift during acquisition was corrected in 3D using a redundant cross-correlation algorithm (Wang et al., 2014) implemented as an independent module of SMAP (Ries, 2020). After translation of the coordinate files, image reconstructions were performed using the ThunderSTORM ImageJ plugin (Ovesný et al., 2014) in Fiji software. Custom scripts and macros were used to translate coordinate files, as well as automate image reconstruction for whole images at 16 nm/pixel for visualization and at 8 nm/pixel for cluster analysis (https://github.com/cleterrier/ChriSTORM).

### PREM

Adherent plasma membranes from rat hippocampal neurons and rat fibroblasts grown on glass coverslips were disrupted by sonication (Heuser, 2000). Unroofed cells were fixed and processed as described previously (Vassilopoulos et al., 2019). Cells were sequentially treated with 0.5% OsO_4_, 1% tannic acid, and 1% uranyl acetate prior to graded ethanol dehydration and hexamethyldisilane substitution (Sigma-Aldrich). Dried samples were rotary-shadowed with 2 nm of platinum and 6 nm of carbon using a high vacuum sputter coater (Leica Microsystems). The resultant platinum-replica was floated off the glass with hydrofluoric acid (5%), washed several times with distilled water, and picked up on 200 mesh formvar/carbon-coated EM grids. Images were captured using a digital camera (Xarosa) and the grids were mounted on an eccentric side-entry goniometer stage of a transmission electron microscope running at 120 kV (Jeol). Images were adjusted for brightness and contrast in Adobe Photoshop (Adobe) and presented in inverted contrast. Tomograms were created by acquiring images with 5° increments at tilt angles up to 20° relative to the sample plane. In Photoshop, layers were used to stack images on top of one another to align them.

### Correlative SIM-PREM/SMLM-PREM microscopy

Unroofed and stained samples were imaged by SIM and an objective-style diamond scriber (Leica) was used to engrave a 1 mm circle around the imaged area. Low-magnification, 10x and 40x phase and fluorescence images were taken as a reference for relocation. After the acquisition session, samples were stored, shipped and platinum replicated as described in the PREM section. After replica generation and prior to its release, the coated surface of each coverslip was scratched with a needle to make an EM grid sized region of interest containing the engraved circle. Grids were imaged with low-magnification EM to relocate the region that was previously imaged by SIM or SMLM, and high magnification EM views were taken from the corresponding axonal region. The SIM or SMLM reconstructions were mapped and aligned by affine transformation to the corresponding high-magnification EM view using the eC-CLEM plugin in ICY software (Paul-Gilloteaux et al., 2017).

### Data quantification and analysis

#### Clathrin pits and dextran clusters segmentation (SIM data)

Clathrin and dextran clusters were segmented using a custom script in Fiji (Schindelin et al., 2012). After manually delineating ROIs corresponding to the proximal axon on a stack of SIM images, fluorescence within each axon ROI was thresholded (threshold levels were determined by the Moments method for clathrin and dextran intact, Otsu method for clathrin and dextran unroofed, eventually fine-tuned manually). Thresholded areas were then segmented using the StarDist plugin in Fiji (Schmidt et al., 2018), with a baseline threshold of 0.5, eventually fine-tuned manually (other StarDist parameters were: StarDist2D Versatile model, normalize bottom 0.1 top 99.8, overlap threshold 0.33). Only the clusters inside the axon ROI or overlapping it by more than half their area were kept, and segmented aggregates of several clusters were removed. The remaining clusters were quantified for area within Fiji and the quantifications exported to a CSV file for each image. The particles CSV files were analyzed in a dedicated JupyterLab notebook, pooling the experiments per condition, and performing particle area, count, density, and pixel-occupation averages.

#### Average image of clathrin pits and dextran clusters

The ROI sets for clusters obtained after the segmentation of SIM images were used to generate average intensity plots of clathrin pits and dextran clusters. 15×15 pixels centered areas were extracted from each ROI and averaged over all ROIs for each channel. The radial average fluorescence profile for each channel was obtained from the averaged image using the Radial Profile Extended ImageJ plugin.

#### Live-cell temporal autocorrelation and kymographs

Live-cell imaging movies were corrected for drift using the Image Stabilization plugin and for bleaching by histogram matching, both in Fiji. Line ROIs were traced on the axon, dendrites, and cell body using NeuronJ (Meijering et al., 2004) on the maximum projection of the movie frames. These line ROIs (360-390 nm line thickness) were used to generate kymographs using the KymoResliceWide plugin (https://github.com/ekatrukha/KymoResliceWide). In addition, the intensity profile along thick line ROIs was used to calculate a temporal correlation profile with the ImageCorrelationJ plugin, comparing the image similarity between all pairs of frames distant by 1, 2, …, (n-1) frames within an n-frames movie (https://www.gcsca.net/IJ/ImageCorrelationJ.html). Temporal correlation profiles were averaged for ROIs of the same type (axon, dendrite, cell body), resulting in an average temporal correlation profile for this compartment.

#### Particle quantification by PREM

Clathrin structures were manually delineated on unroofed PM pictures using ImageJ. Each object was classified as a CCP by the observer and the object area was measured. On each image, the total membrane area was also measured to calculate the object density (object count normalized by membrane area).

### Statistics

Individual measurement points (n) from independent experiments (N) were pooled. Intensity profiles, graphs, and statistical analyses were generated using Prism. On bar graphs, dots (if present) are averages of each independent experiment, bars or horizontal lines represent the mean, and vertical lines are the SEM unless otherwise specified. Significances were tested using one-way, non-parametric ANOVA with Kruskal-Wallis posthoc significance testing between selected conditions. In Figures, the results of the post-hoc significance are indicated as follows: ns or ns, non-significant; ^*^, p < 0.05; ^**^, p < 0.01; ^***^, p < 0.001.

## Supporting Material

**Supporting Figure 1:**
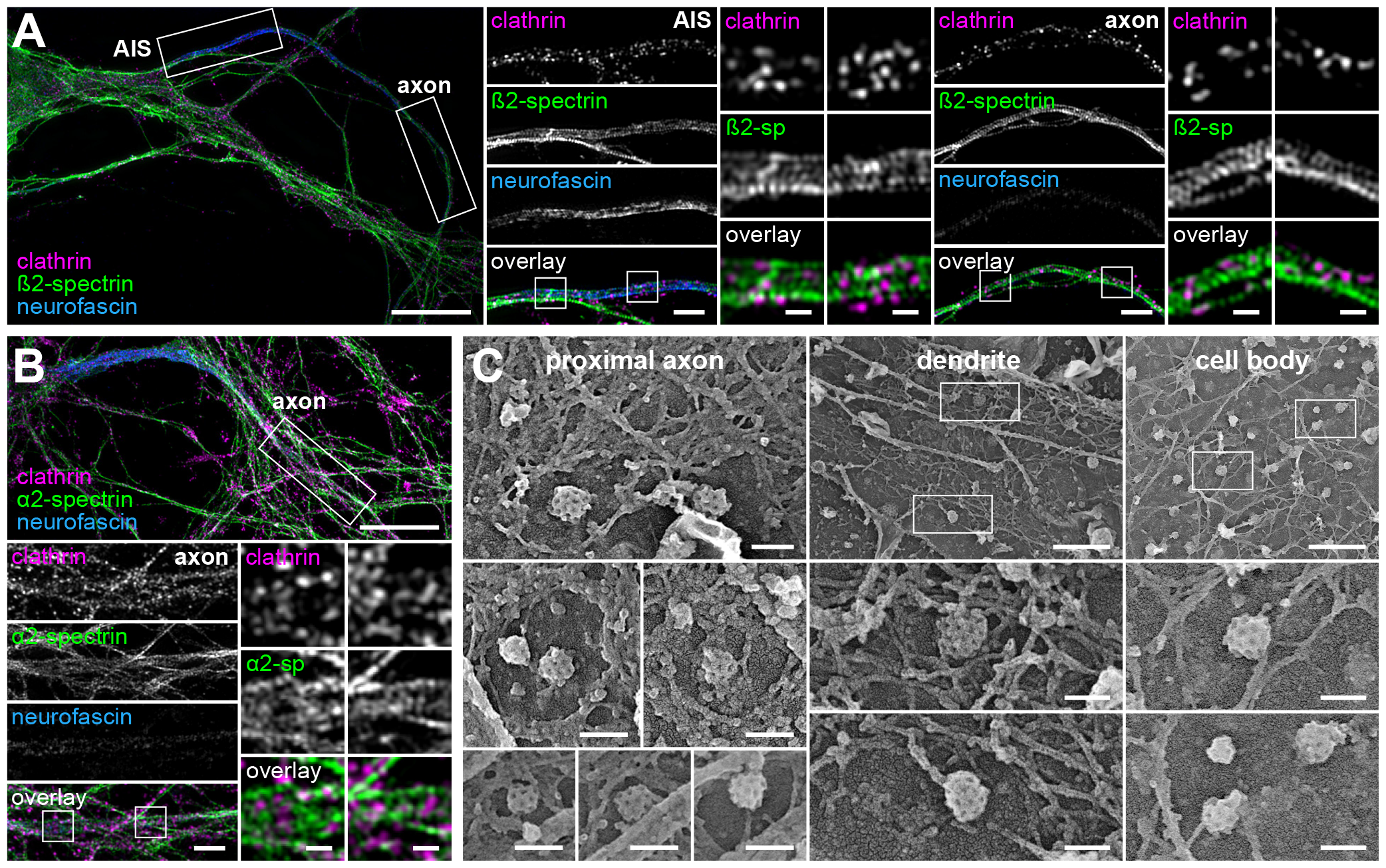
Clearings of the periodic spectrin mesh encase CCPs along the AIS and proximal axon, but not in the dendrites or cell body of neurons. A. SIM image of a cultured hippocampal neuron at 14 div fixed and stained for clathrin (magenta, pits appear as clusters), ß2-spectrin (green, ∼190 nm-spaced bands are visible) and neurofascin (blue, labels the AIS). Zooms show portions of the AIS (center panels) and axon proper after the AIS (right panels). Scale bars, 10 µm (left image), 2 µm (center column), 0.5 µm (square zoomed images). B. Same SIM image as in Fig. 1A of a cultured hippocampal neuron at 14 div fixed and stained for clathrin (magenta), α2-spectrin (green) and neurofascin (blue), with a zoom on the axon proper (bottom panels). Scale bars, 10 µm (top image), 2 µm (bottom left panels), 0.5 µm (bottom right square zoomed images). C. Left, PREM views showing CCPs found along the proximal axon in circular areas devoid of the actin-spectrin scaffold (left), or along dendrites (center) and in the cell body (right) where the submembrane scaffold is sparser. Scale bars, 100 nm.

**Supporting Figure 2:**
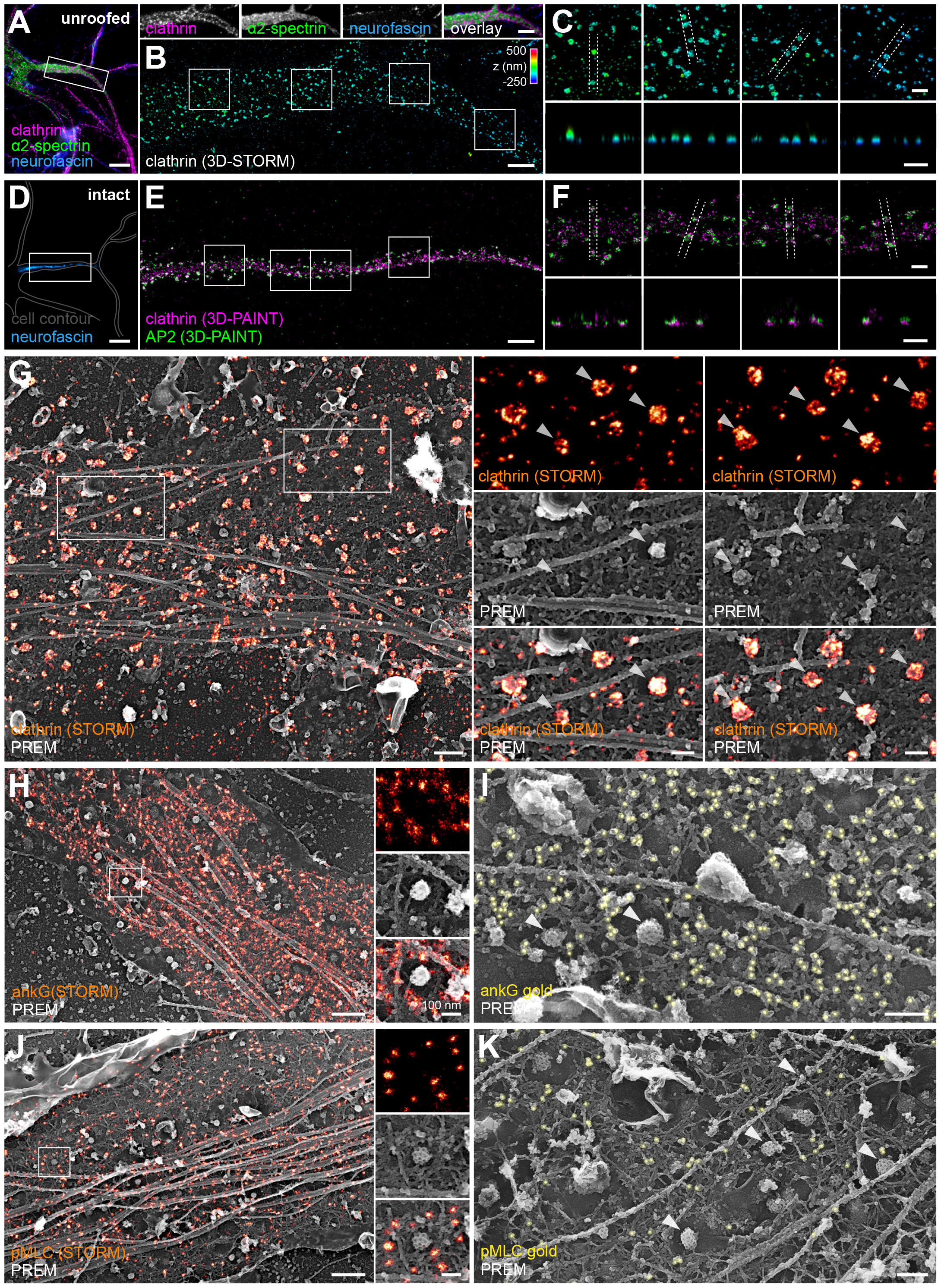
Molecular organization of CCPs, ankyrin G and pMLC along AIS. (next page >) A. Widefield image of a neuron unroofed, fixed and stained for clathrin (magenta), α2-spectrin (green) and neurofascin (blue). Line of images on the right are individual channels and overlays corresponding to the AIS area highlighted on the left image. Scale bars, 10 µm (left image), 5 µm (AIS images). B. 3D-STORM image of clathrin corresponding to the AIS area shown in A, color coded for depth. Scale bar, 2 µm. C. Zooms showing individual CCPs on XY images (top row, corresponding to areas highlighted in B) and corresponding XZ transverse sections (bottom row, taken between the lines highlighted on the XY images). Pits are found along the remaining ventral plasma membrane after unroofing. Scale bars, 0.5 µm. D. Widefield image of a neuron fixed and stained for neurofascin (blue). The contour of the whole neuron is delineated (gray). Scale bar, 10 µm. E. 2-color 3D-PAINT images of the AIS corresponding to the image in D with staining for clathrin (magenta) and AP2 (green). Scale bar, 2 µm. F. Zooms showing individual CCPs immunolabeled for clathrin heavy chain (magenta) and AP2 (green) on XY images (top row, corresponding to areas highlighted in E) and corresponding XZ transverse sections (bottom row, taken between the lines highlighted on the XY images). Both CHC and AP2 colocalize at CCPs. Scale bars, 0.5 µm. G. Correlative STORM-PREM image of an unroofed AIS immunolabeled for clathrin (orange). Right, zoomed images showing CCPs in spectrin mesh clearings. Scale bars, 500 nm (left image), 100 nm (zoomed images). H. Correlative STORM-PREM image of an unroofed AIS immunolabeled for ankyrin G (ankG, orange). Right, zoomed images showing CCPs in spectrin mesh clearings. Scale bars, 500 nm (left image), 100 nm (zoomed images). I. PREM view of an unroofed AIS immunogold-labeled for ß4-spectrin (orange), showing CCPs in spectrin mesh clearings, with 15 nm gold beads bound to the spectrin mesh around the bare membrane area. Scale bar, 200 nm (zoomed images). J. Correlative STORM-PREM image of an unroofed AIS labeled for pMLC (orange). Right, zoomed images showing CCPs in spectrin mesh clearings. Scale bars, 5 µm (left image), 100 nm (zoomed images). K. PREM view of an unroofed AIS immunogold-labeled for pMLC (yellow), showing CCPs in spectrin mesh clearings, with 15 nm gold beads bound to the spectrin mesh around the bare membrane area. Scale bar, 200 nm.

**Supporting Figure 3:**
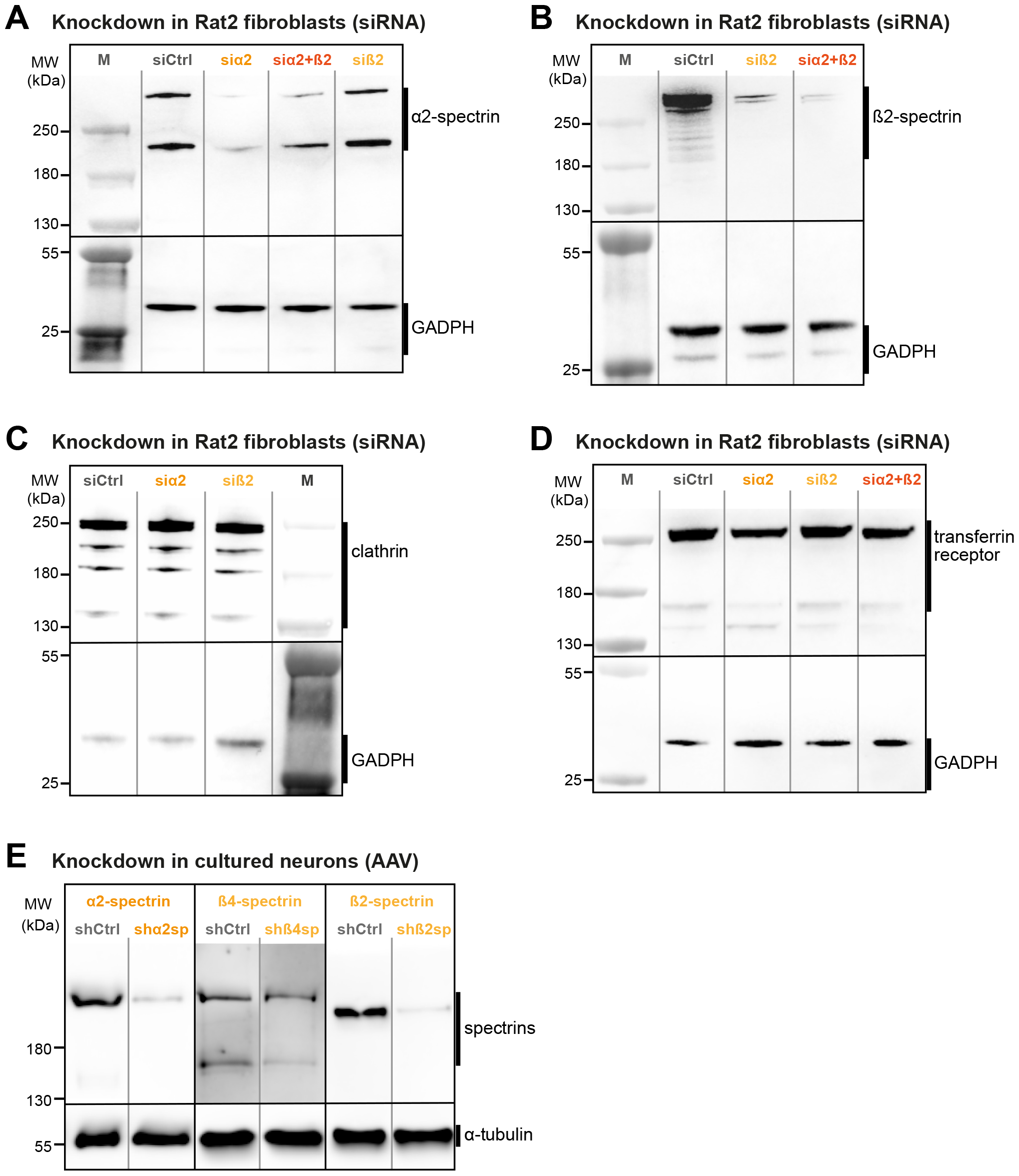
siRNA and shRNA knockdown efficiency assessed by Western blot. A. Western blot of lysates from Rat2 fibroblasts transfected with control, α2-spectrin, α2+ß2-spectrin, or ß2-spectrin siRNAs probed for α2-spectrin (top), with GADPH as a loading control (bottom). B. Western blot of lysates from Rat2 fibroblasts transfected with control, ß2-spectrin, or α2+ß2-spectrin siRNAs probed for ß2-spectrin (top), with GADPH as a loading control (bottom). C. Western blot of lysates from Rat2 fibroblasts transfected with control, α2-spectrin, or ß2-spectrin siRNAs probed for clathrin (top), with GADPH as a loading control (bottom). Clathrin levels are unaffected by the knockdown of α2 and/or ß2-spectrin. D. Western blot of lysates from Rat2 fibroblasts transfected with control, α2-spectrin, ß2-spectrin, or α2+ß2-spectrin siRNAs probed for transferrin receptor (TfR, top), with GADPH as a loading control (bottom). TfR levels are unaffected by the knockdown of α2 or ß2-spectrin. E. Western blots of lysates from hippocampal neurons infected with control, α2-spectrin, ß4-spectrin, or ß2-spectrin shRNA AAVs probed for α2-spectrin, ß4-spectrin, or ß2-spectrin, respectively. α-tubulin is used as a loading control (bottom).

**Supporting Figure 4:**
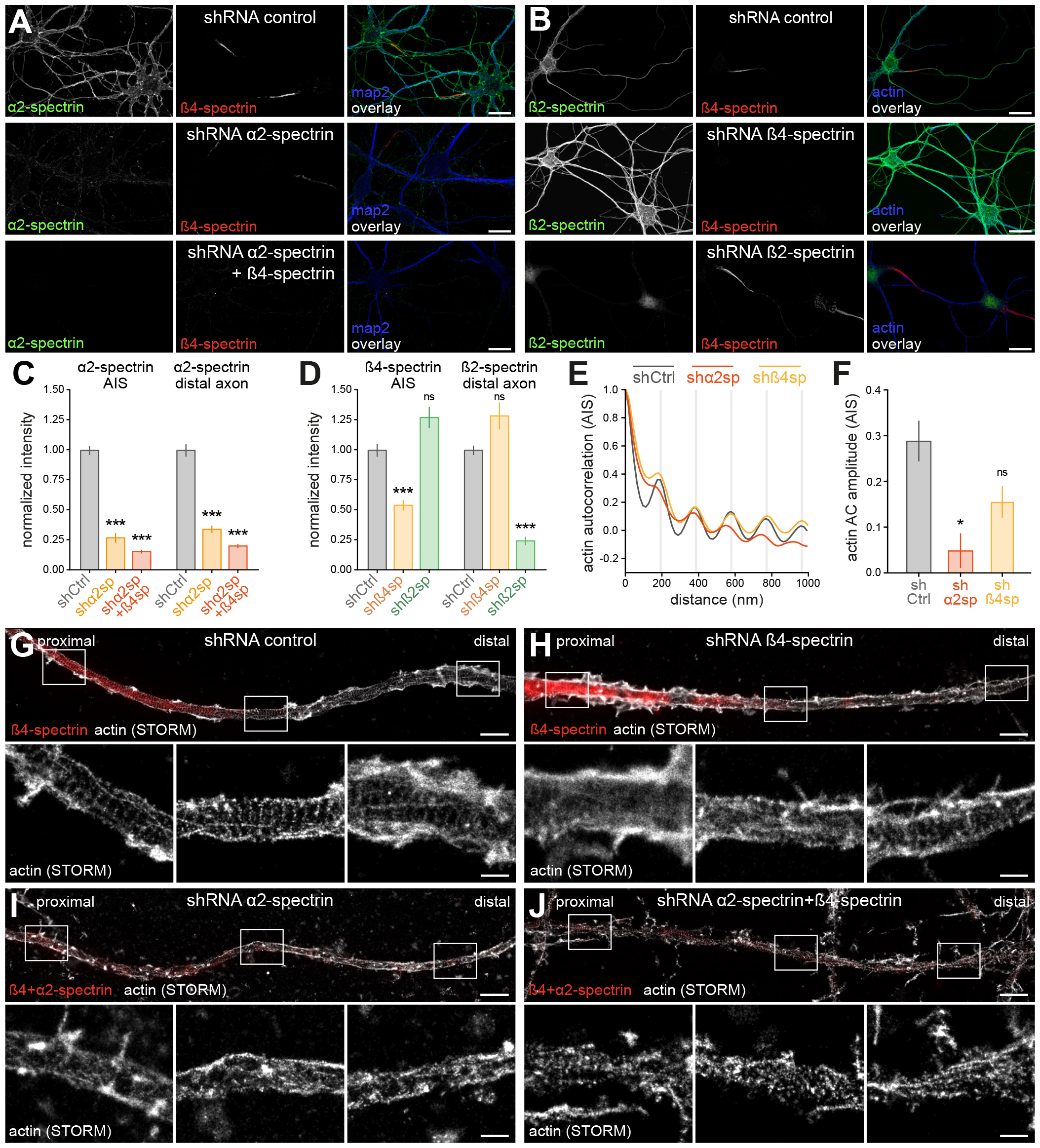
Effect of spectrin knockdown in neurons assessed by immunolabeling and STORM. A. Epifluorescence images of cultured neurons infected with control, α2-spectrin, or α2+ß4-spectrin shRNA AAVs, fixed and stained for α2-spectrin (green), ß4-spectrin (red) and map2 (blue). Scale bars, 20 µm. B. Epifluorescence images of cultured neurons infected with control, ß4-spectrin, or ß2-spectrin shRNA AAVs, fixed and stained for ß2-spectrin (green), ß4-spectrin (red) and map2 (blue). Scale bars, 20 µm. C. Quantification of the normalized staining intensity for α2-spectrin at the AIS (left) and distal axon (right) in neurons infected with control, α2-spectrin, or α2+ß4-spectrin shRNA AAVs, showing the efficient knockdown of α2-spectrin in both compartments. D. Quantification of the normalized staining intensity for ß4-spectrin at the AIS (left) and ß2-spectrin in the distal axon (right) in neurons infected with control, ß4-spectrin, or ß2-spectrin shRNA AAVs, showing the selective knockdown of each ß-spectrin in the AIS and distal axon. E. Autocorrelation curves from intensity profiles traced along the AIS of neurons infected with control, α2-spectrin, or ß4-spectrin shRNA AAVs, fixed, stained for actin using phalloidin-Alexa Fluor 647, and imaged by STORM (see G-J). Oscillations demonstrating the 190-nm periodicity (actin rings) are attenuated in α2-spectrin and ß4-spectrin knocked-down neurons. F. Quantification of the autocorrelation amplitude (first valley to first peak) on the graph in E, showing lower values in α2-spectrin and ß4-spectrin knocked-down neurons. G. Image of the AIS of a neuron infected with a control shRNA AAV, fixed, and stained for ß4-spectrin (red, epifluorescence image) and actin (gray, STORM image). Bottom zooms show the actin rings in the proximal, middle, and distal portions of the axon shown. Scale bar, 500 nm. H. Image of the proximal axon of a neuron infected with a ß4-spectrin shRNA AAV, fixed, and stained for ß4-spectrin (red, epifluorescence image) and actin (gray, STORM image). Bottom zooms show the selective perturbation of the actin rings in the proximal portion of the axon shown. Scale bar, 500 nm. I. Image of the proximal axon of a neuron infected with a α2-spectrin shRNA AAV, fixed, and stained for α2+ß4-spectrin (red, epifluorescence image) and actin (gray, STORM image). Bottom zooms show the perturbation of the actin rings in the proximal, middle, and distal portions of the axon shown. Scale bar, 500 nm. J. Image of the proximal axon of a neuron infected with an α2+ß4-spectrin shRNA AAV, fixed, and stained for α2+ß4-spectrin (red, epifluorescence image) and actin (gray, STORM image). Bottom zooms show the perturbation of the actin rings in the proximal, middle, and distal portions of the axon shown. Scale bar, 500 nm.

**Supporting Movie 1:**
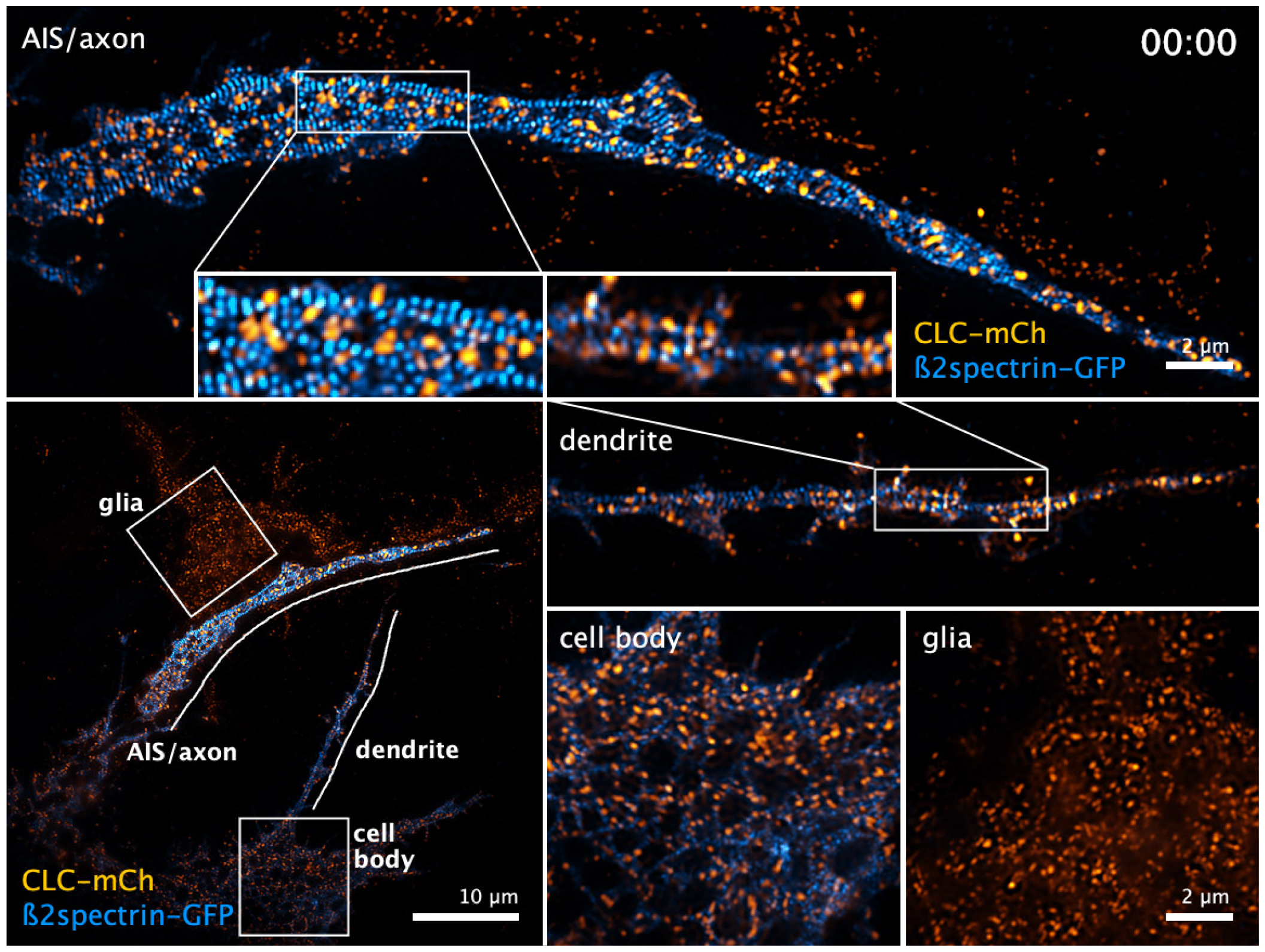
Dynamics of neuronal CCPs and the axonal periodic scaffold by live-cell TIRF-SIM. Hippocampal neuron in culture transfected at 10 div with clathrin light chain-mCherry (CLC-mCh, orange) and EGFP-ß2-spectrin (ß2spectrin-GFP, blue) and imaged at 13 div by 2-color TIRF-SIM (one frame every 20s for 20 min, 61 frames total). Reconstructed TIRF-SIM images were denoised using a supervised deep-learning model trained on pairs of low and high SNR images of fixed cells expressing the same constructs (Enhance.ai). Bottom left, full-field view of the transfected neuron with highlighted areas corresponding to the AIS/proximal axon (top), dendrite (middle right), cell body (bottom center), and a neighboring glia only transfected with CLC-mCh (bottom left). Further zooms detail the dynamic of CCPs within the AIS and along the dendrite. CCPs are more stable at the AIS, being present throughout the 20-minute movie, whereas they appear more dynamic in the dendrites and cell body, and very dynamic in the glial cell.

**Supporting Movie 2:**
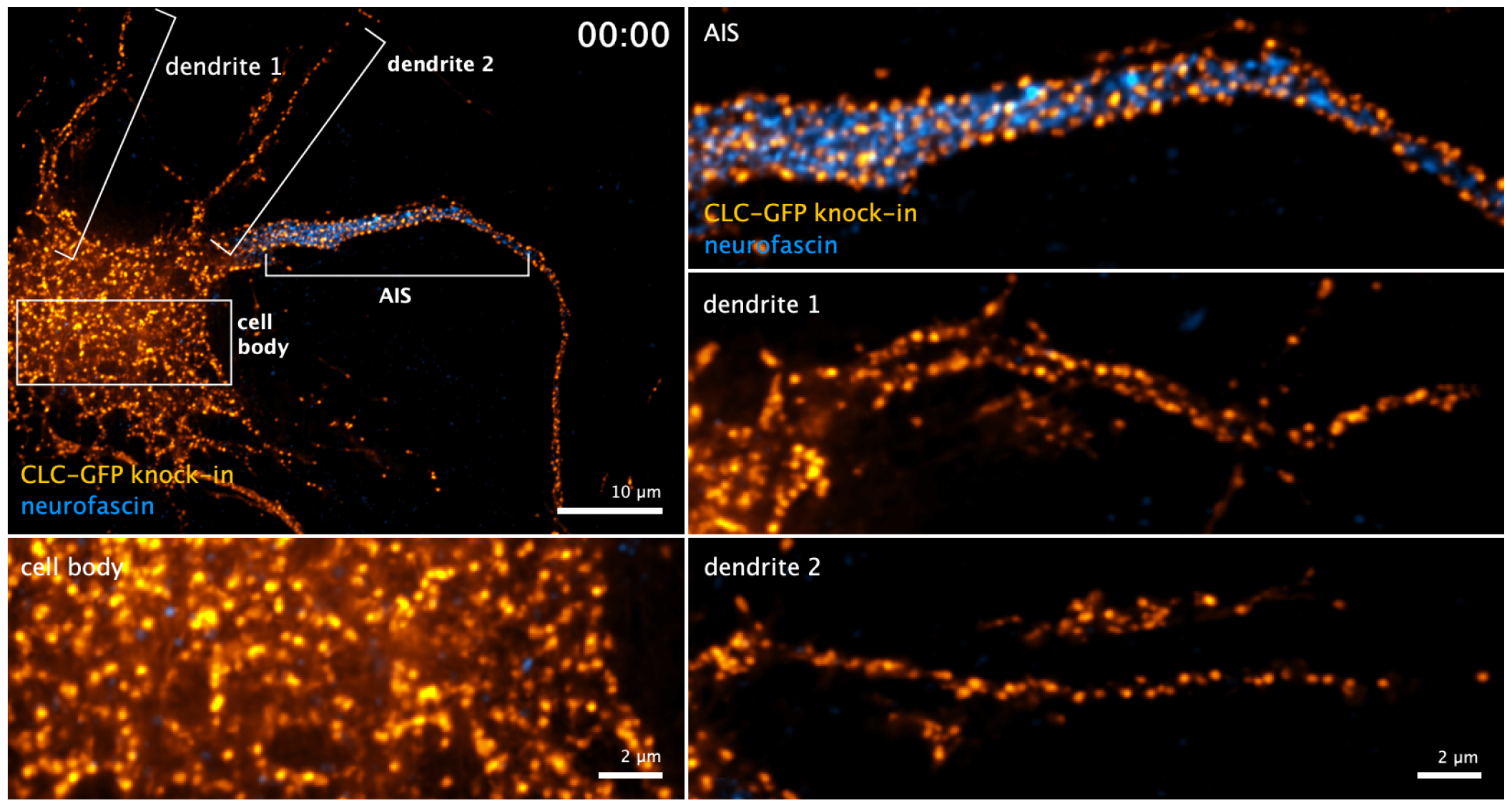
Dynamics of neuronal CCPs using super-resolved imaging of endogenously-tagged clathrin. Hippocampal neuron in culture infected with AAVs targeting the endogenous tagging of clathrin light chain with EGFP at its C-terminus (CLC-GFP knock-in, orange), labeled for the AIS using an anti-neurofascin antibody targeting an extracellular epitope (blue), and imaged at 14 div using SoRa super-resolved spinning disk microscopy (one frame every 5s for 10 m, 121 images total). Raw images were denoised using a built-in deep-learning module trained on a dataset of confocal images (Denoise.ai). Top left, full-field view of the transfected neuron with highlighted areas corresponding to the AIS/proximal axon (top right), two dendrites (middle right and bottom right), and the cell body (bottom left). CCPs are more stable at the AIS compared to the dendrites and cell body.

## Notes

### Competing Interest Statement

The authors have declared no competing interest.

